# Identification of a novel missense variant in *SPDL1* associated with idiopathic pulmonary fibrosis

**DOI:** 10.1101/2020.06.29.178079

**Authors:** Ryan S. Dhindsa, Johan Mattsson, Abhishek Nag, Quanli Wang, Louise V. Wain, Richard Allen, Eleanor M. Wigmore, Kristina Ibanez, Dimitrios Vitsios, Sri VV. Deevi, Sebastian Wasilewski, Maria Karlsson, Glenda Lassi, Henric Olsson, Daniel Muthas, Alex Mackay, Lynne Murray, Simon Young, Carolina Haefliger, FinnGen Consortium, Toby M. Maher, Maria G. Belvisi, Gisli Jenkins, Philip Molyneaux, Adam Platt, Slavé Petrovski

## Abstract

Idiopathic pulmonary fibrosis (IPF) is a fatal disorder characterised by progressive, destructive lung scarring. Despite significant progress, the genetic determinants of this disease remain incompletely defined. Using next generation sequencing data from 752 individuals with sporadic IPF and 119,055 controls, we performed both variant- and gene-level analyses to identify novel IPF genetic risk factors. Our variant-level analysis revealed a novel rare missense variant in *SPDL1* (NM_017785.5 p.Arg20Gln; *p* = 2.4 × 10^−7^, odds ratio = 2.87). This signal was independently replicated in the FinnGen cohort (combined *p* = 2.2 × 10^−20^), firmly associating this variant as a novel IPF risk allele. *SPDL1* encodes Spindly, a protein involved in mitotic checkpoint signalling during cell division that has not been previously described in fibrosis. Our results highlight a novel mechanism underlying IPF, providing the potential for new therapeutic discoveries in a disease of great unmet need.

## Main

Idiopathic pulmonary fibrosis (IPF) is a progressive scarring disorder of the lung that preferentially affects individuals over the age of 70^1^. In the absence of a lung transplant, individuals with IPF have an average life expectancy of three to five years after diagnosis^2^. Approved drugs are not curative and are associated with considerable side effects making them poorly tolerated^3^. Identifying genetic risk factors of IPF helps elucidate disease aetiology, which is crucial to the development of more precise therapies. Furthermore, an improved understanding of genetic risk factors associated with IPF may enable stratification of patients in clinical trials^3^. Genome-wide association studies (GWAS) have implicated common variants at several loci^4^, with the strongest signal mapping to the promoter region of *MUC5B*^5^. Nonetheless, common variants seem to explain a small proportion of IPF heritability compared to rare deleterious variants in the protein-coding region of the genome^6^. Sequencing-based case-control studies have consistently identified three definitive IPF risk genes in both familial and sporadic forms of IPF: *TERT*, *RTEL1*, and *PARN*, all involved in telomerase biology^6,7^. Rare variants in the telomerase RNA component, *TERC*, have also been implicated in sporadic IPF^7^. Despite the significant progress in identifying both rare and common variant signals, the underlying genetic predisposition remains unknown for the majority of IPF patients.

We performed variant- and gene-level analyses to identify novel IPF risk factors using genetic sequencing data from 752 European cases with IPF and 119,055 European controls (**Extended Data Fig. 1a,b**). These 752 cases specifically comprised 507 individuals enrolled in PROFILE (Prospective Study of Fibrosis In the Lung Endpoints) and 245 UK Biobank participants with IPF (ICD10 code J84*) registered as the primary (Field 40001) or secondary (Field 40002) cause of death (**Fig. 1**). The median age of diagnosis for the cases was 71 years of age with a median survival of 39.4 months. The control cohort consisted of UK Biobank participants screened for non-respiratory disease (**Table S1**). Whole-genome sequencing was performed on PROFILE participants and whole-exome sequencing was performed on UK Biobank participants. We therefore limited our association analyses to pruned protein-coding sequence sites with minimum variability in coverage between cases and controls (**Table S2**). Variant-level associations were replicated in the FinnGen cohort, which includes genotype data for individuals of Finnish descent, including 1,028 individuals with IPF and 196,986 controls (FinnGen release 5) (**Fig. 1**). We also performed a combined gene-level collapsing analysis using data from a previously published whole-exome sequencing study that included 262 cases and 4,141 controls (**Fig. 1**).

**Figure 1.**
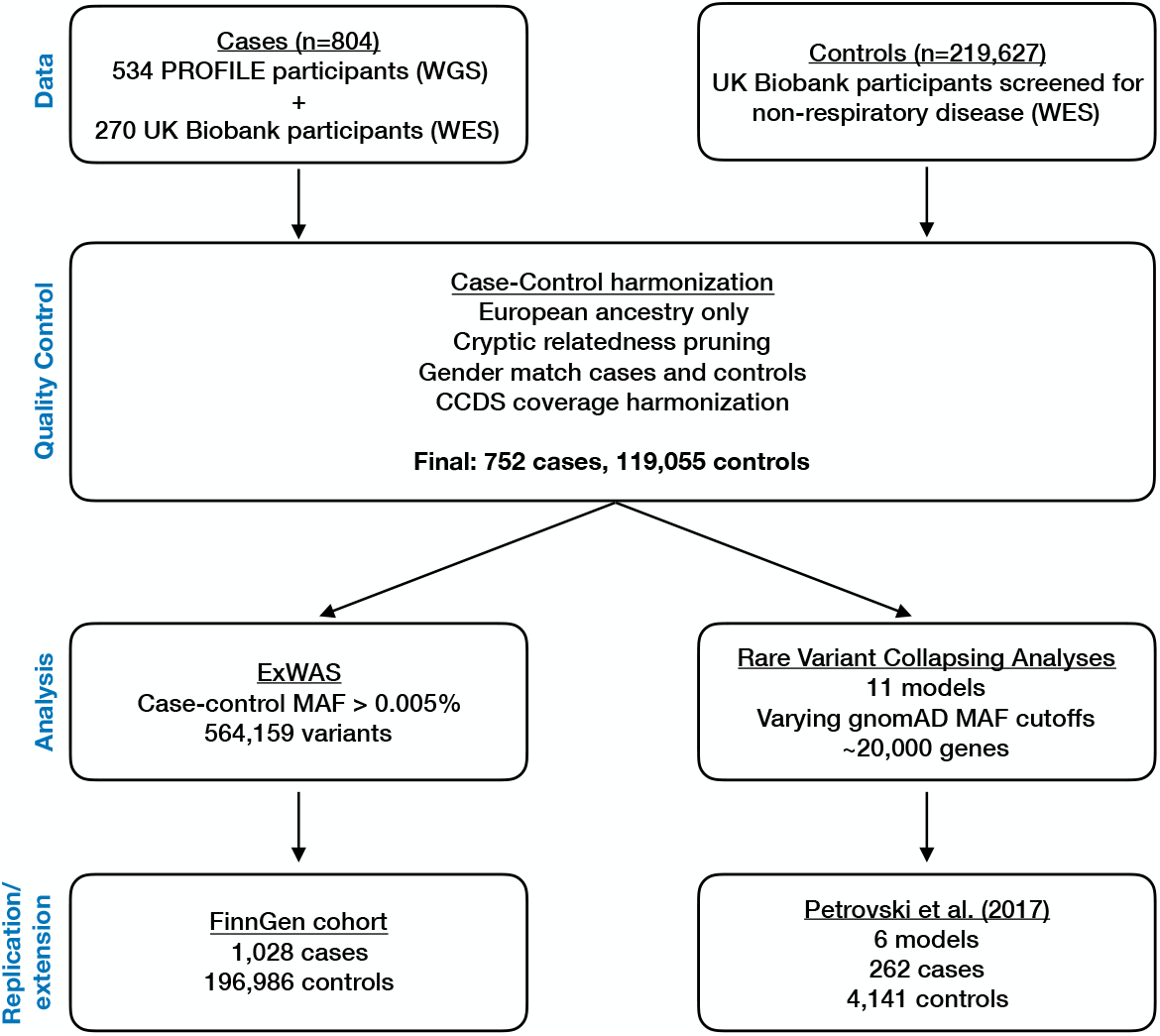
Idiopathic pulmonary fibrosis genetic discovery and replication study design. We combined whole-exome sequence (WES) and whole-genome sequence (WGS) data from a total of 804 cases enrolled in either the PROFILE study or the UK Biobank. Controls comprised of 219,627 UK Biobank participants. We harmonized the case-control cohort based on ancestry, relatedness, and gender, resulting in a total of 752 cases and 119,055 controls screened for non-respiratory disease. Furthermore, we filtered out sites that were differentially covered between cases and controls. We then performed two analyses: an exome-wide variant-level association test (ExWAS) and rare variant collapsing analysis. Suggestive variants from the ExWAS were then replicated in the FinnGen cohort. Collapsing analyses were also combined with previously published results from an independent case-control study^6^.

**Figure 2.**
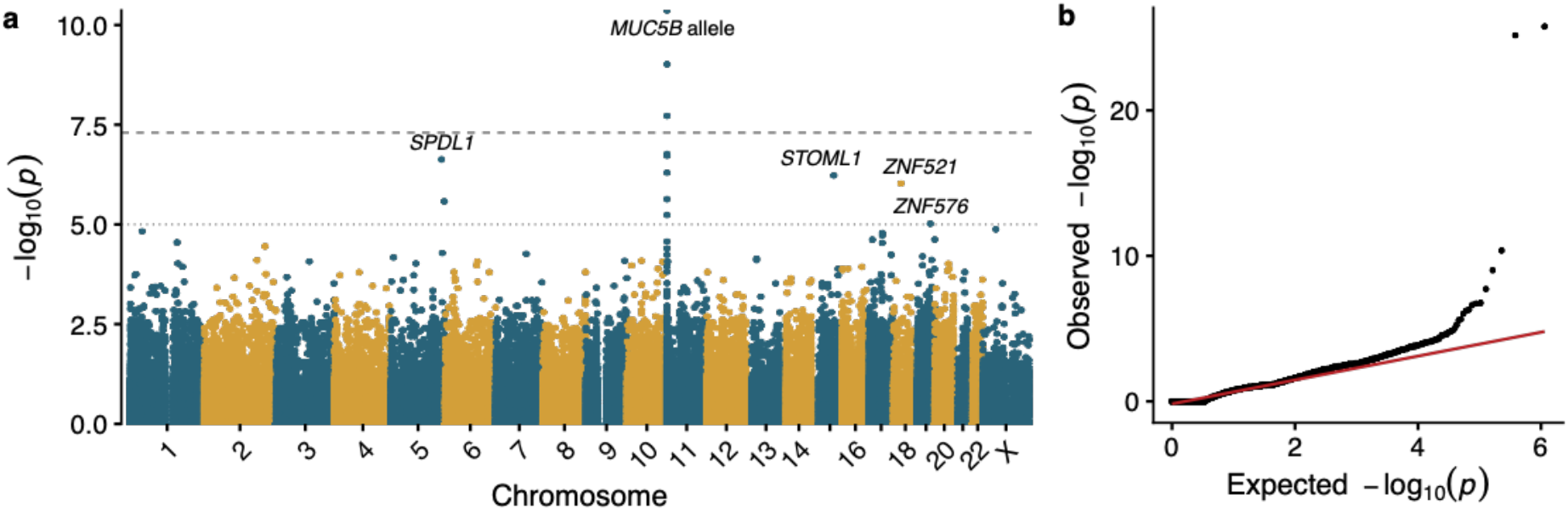
Association of single-nucleotide variants with IPF. **(a)** Manhattan plot depicting *p*-values of the 564,159 exonic variants tested for association with IPF status. The long-dash line indicates the genome-wide significance threshold (p < 5 × 10-8). Relevant to the *MUC5B* locus, the Y-axis is capped at 1×10^−10^. **(b)** Quantile-quantile plot of observed versus expected *p*-values.

In an exome-wide variant-level association study (exWAS), we assessed 564,159 protein-coding variants for association with IPF risk. We identified five genome-wide-significant variants (*p* < 5 × 10^−8^), all in the vicinity of the well-established *MUC5B* risk allele (rs35705950). The next strongest independent signal emerged from a missense variant in the gene *SPDL1* (NM_017785.5 p.Arg20Gln [rs116483731]; *p* = 2.35 × 10^−7^), with an allele frequency of 2.2% in cases compared to 0.78% in controls (odds ratio [OR], 2.87). This same variant reached genome-wide significance in the FinnGen replication cohort (**Extended Data Fig. 2**), with a case frequency of 6.9% and a control frequency of 3.0% (*p* = 9.97 × 10^−16^; OR, 3.13). Taking the combined evidence from both cohorts, this variant achieves a Stouffer Z-test *p*-value of 2.2 × 10^−20^, unequivocally associating it with IPF risk.

Despite its relatively strong effect size, the *SPDL1* locus has not been previously reported in IPF through prior GWAS with larger sample sizes (**Table S3**)^4^. Because of known Mendelian genetics contribution to IPF genetic architecture, we tested whether the *SPDL1* missense variant may have independently arisen multiple times in Europeans or whether it resides on a common haplotype. We examined the haplotypes in the 10kb window of the *SPDL1* index variant and found that all *SPDL1* rs116483731 risk allele observations among the PROFILE cohort occurred on a single common ancestral haplotype that accounted for 16.5% of all haplotypes identified among the 1,014 PROFILE chromosomes (**Extended Data Fig. 3**). This indicates a common ancestral origin for this variant.

We next performed gene-based collapsing analyses to identify genes carrying an aggregated excess of rare deleterious variants among the case sample. Despite this being the largest gene-based collapsing test performed in IPF to date, no new genes reached study-wide significance (*p* < 2.4 × 10^−7^) across 11 different rare-variant genetic architectures (**Extended Data Fig. 4; Table S5**). We then combined our results with data reported in a previously published study of 262 IPF cases and 4,141 controls^6^, which also did not yield novel study-wide significant findings (**Extended Data Fig. 5** and **Table S6**). In addition, we explored whether the top-ranked genes that did not meet study-wide significance were enriched for novel, putative disease-associated genes. Using mantis-ml^8^, we found that case-enriched genes (p < 0.05) in our collapsing models that focused on variants in regions intolerant to missense variation^9^ were significantly enriched for genes predicted to be associated with pulmonary fibrosis (**Extended Data Fig. 6**). This result suggests that there are additional IPF risk genes to be discovered in larger case sample sizes.

Our primary collapsing model focused on rare (MAF < 0.1%) protein-truncating venric ariants (PTVs). Under this model, the top three genes were the previously reported *RTEL1* (*p* = 3.0 × 10^−7^; OR, 13.6), *PARN* (*p* = 2.1 × 10^−5^; OR, 28.9), and *TERT* (*p* = 8.5 × 10^−5^; OR, 43.3) signals. Given the rarity of this extreme class of variants among these genes in the general population, the effect size of carrying a PTV in these genes came with larger effect sizes than the more common variants implicated in IPF (**Fig. 3a**). Notably, the *MUC5B* and *SPDL1* SNPs represented the next strongest risk factors (**Fig. 3a**).

**Figure 3.**
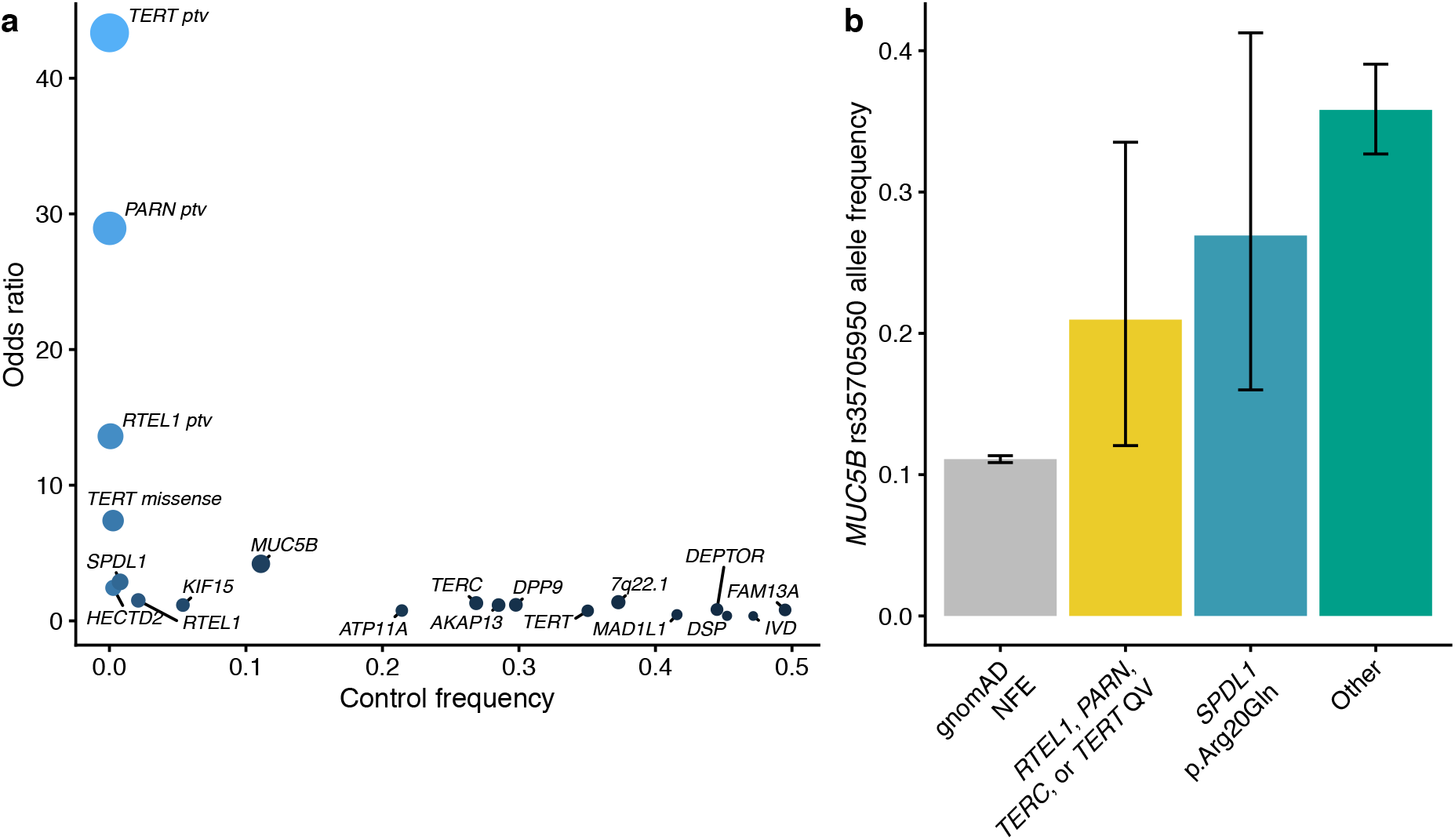
Loci associated with IPF. **(a)** Scatter plot depicting the odds ratios versus control frequencies of protein-truncating variants (PTVs) in *TERT*, *PARN*, and *RTEL1,* rare damaging missense variants in *TERT* based on PROFILE collapsing analyses, the novel missense variant in *SPDL1*, and the sentinel SNPs from the largest IPF GWAS, to date. Control frequencies for *TERT*, *PARN*, and *SPDL1* missense reflect the carrier frequencies in the UK Biobank controls used for our association studies. Control frequencies for the remaining alleles were derived from gnomAD non-Finnish European allele frequencies. **(b)** Allele frequency of the *MUC5B* promoter allele among non-Finnish European gnomAD samples and PROFILE IPF cases stratified by genotype. “Other” refers to IPF cases who do not carry a rare variant in *RTEL1*, *PARN*, *TERC*, *TERT*, and do not carry the *SPDL1* missense variant (i.e. noncarriers).

It is well-established that the *MUC5B* promoter risk allele frequency is significantly enriched in cases carrying rare variants in *RTEL1, TERT,* and *PARN* compared to controls, albeit at a lower rate than among noncarrier cases^6,10^. Using the WGS data available for the cases in the PROFILE cohort, we found that the *MUC5B* allele frequency in carriers of rare variants in *RTEL1, PARN, TERT*, and *TERC* (21%) was significantly higher than the allele frequency in non-Finnish European controls (11%; OR = 2.13, p = 0.02), but lower than IPF cases without an identified rare genetic risk factor in a telomere-related gene (33%; p = 0.02) (**Fig. 3b**). As previously suggested^6^, these results underscore the contribution of an oligogenic architecture contributing to IPF risk.

Both the presence of mutations in telomerase genes such as *TERT*, *TERC, RTEL1* and *PARN* and quantifiably shorter telomere lengths have been associated with poorer prognosis in individuals with IPF^11^. Given this, we used TelSeq^12^ to infer the telomere lengths from the WGS data for the 507 cases included in the PROFILE cohort. We then tested whether different genetic risk factors were associated with telomere length using a logistic-regression model that included sex and age as covariates. Indeed, the telomeres of PROFILE IPF cases carrying rare putatively pathogenic variants in *RTEL1*, *PARN*, *TERC* or *TERT* were 10-15% shorter than the telomeres of the remaining cases in the PROFILE cohort (*p* = 0.004) **(Fig. 4**; **Table S7, S8**). In keeping with prior reports, individuals with these variants were gender balanced (in contrast to the wider PROFILE cohort, which was three quarters male), younger, and had worse survival rates than that seen for the cohort as a whole (**Table S7 and Extended Data Fig. 7a-c**). Individuals carrying the minor allele for *MUC5B* or *SPDL1* did not exhibit statistically significant differences in their telomere lengths compared to the remainder of the PROFILE cohort **(Fig. 4**) nor did they exhibit significant differences in demographic or clinical characteristics (**Table S7**). There was no significant enrichment of family history among individuals carrying rare variants in the telomerase genes, the *SPDL1* risk allele, or the *MUC5B* risk allele (**Table S7**).

**Figure 4.**
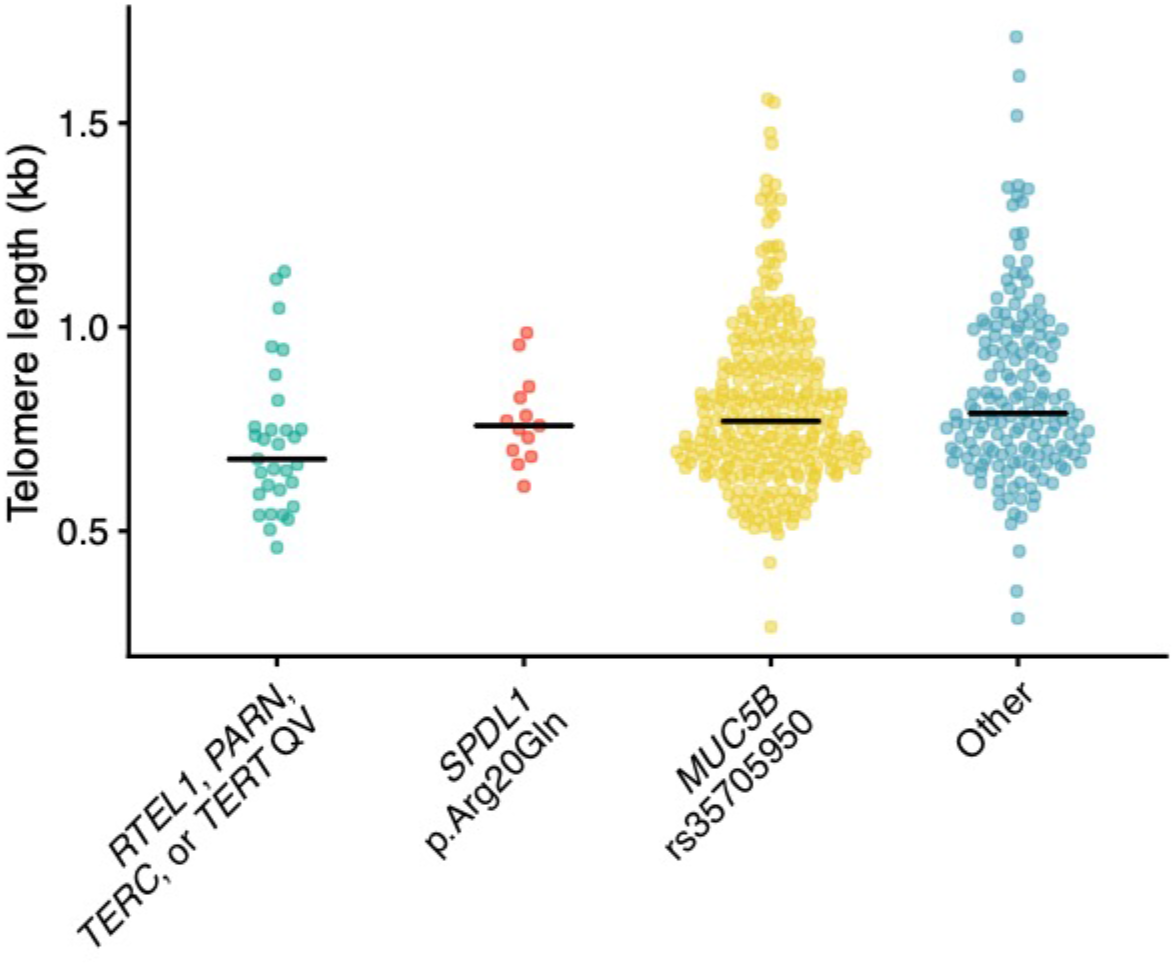
Telomere lengths in individuals with IPF. TelSeq-inferred^12^ telomere lengths for the 507 IPF cases included in the PROFILE cohort stratified by genetic risk factors.

The differences in telomere lengths may be partially explained by gene function. *RTEL1*, *PARN*, and *TERC,* and *TERT* are all directly involved in the telomerase pathway, whereas *SPDL1* encodes spindly, a coiled-coil domain containing protein that has a critical role in mitosis. This protein regulates chromosome alignment as well as microtubule attachment to kinetochores during prometaphase^13–15^. It also regulates the spindle assembly checkpoint (SAC) and enables kinetochore compaction by recruiting the microtubule motor dynein to kinetochores, facilitating the removal of outer kinetochore components and SAC proteins. Following the formation of stable microtubule attachments, these processes allow cells to progress from metaphase into anaphase and to complete mitosis. Mechanistically, Spindly functions as an adaptor protein, linking the RZZ complex (Rod, Zw10 and Zwilch) with dynein/dynactin. Spindly binds dynein and dynactin via two C-terminal domains termed the CC1 box and the Spindly Motif respectively. Importantly, cells carrying mutant forms of Spindly lacking these domains have been shown to display aberrant spindle morphology and chromosome segregation errors^14,15^. In addition to its role in mitosis, Spindly has also been reported to localise to the leading edge of migrating cells^16^. Cells either knocked down for Spindly or expressing a mutant version that is deficient in dynactin binding showed reduced migration in a wound scratch assay. This mechanism has been suggested to be involved in the spread of colorectal cancer cells^17^. Recently, two other genes also linked to kinetochore function, *MAD1L1* and *KIF15*, were implicated in an IPF GWAS study^4^, suggesting that dysfunction of this pathway may underlie a novel non-telomeric mechanism in IPF. Individuals with the *SPDL1* minor allele, at least superficially, resemble the wider cohort of IPF subjects (**Table S7**). Deeper phenotyping of PROFILE and other IPF cohorts may uncover unique features of disease related to impaired kinetochore function. It will also be important to explore whether genetic mechanisms influence response to anti-fibrotic therapy—a finding which would influence future trial design and precision medicine strategies. Notably, only one individual in the case cohort carried both the *SPDL1* minor allele and a qualifying variation in either *RTEL1*, *PARN*, *TERC,* or *TERT*, further suggesting that these are two independent mechanisms of IPF pathogenesis, and that mitotic spindle dysfunction pathway underlies a novel non-telomeric mechanism in IPF.

In conclusion, our work unveiled a novel genetic risk factor for IPF. Our results also underscore the advantage of sequencing-based variant-association tests in capturing signals across the allele frequency spectrum. Furthermore, the higher frequency of this variant in the Finnish population highlights the value of performing genetic analyses in bottleneck populations. Although one might speculate of a role for Spindly in cell senescence or fibroblast migration in pulmonary fibrosis, this finding necessitates further experimental follow-up. Importantly, the evidence of multiple pathways for this devastating disorder also emphasizes the need for more targeted treatments.

## Methods

### Sequencing, alignment, and variant calling

For the PROFILE cohort, genomic DNA from IPF cases was extracted and underwent paired-end 150bp WGS at Human Longevity Inc using the NovaSeq6000 platform. For IPF cases, >98% of consensus coding sequence release 22 (CCDS) has at least 10x coverage and average coverage of the CCDS achieved 42-fold read-depth. Genomic DNA from UK Biobank controls underwent paired-end 75bp whole exome sequencing (WES) at Regeneron Pharmaceuticals using the IDT xGen v1 capture kit on the NovaSeq6000 machines. For UK Biobank controls, >95% of CCDS has at least 10x coverage and average CCDS read-depth of 59X. All case and control sequences were processed through the same bioinformatics pipeline, this included re-processing all the UK Biobank exomes from their unaligned FASTQ state. A custom-built Amazon Web Services (AWS) cloud compute platform running Illumina DRAGEN Bio-IT Platform Germline Pipeline v3.0.7 was adopted to align the reads to the GRCh38 genome reference and perform small variant SNV and indel calling. SNVs and indels were annotated using SnpEFF v4.3 against Ensembl Build 38.92.

### Ethics statement

For the PROFILE cohort written informed consent was obtained from all subjects and the study was approved by the local research ethics committee (reference numbers 10/H0720/12).

Patients and control subjects in FinnGen provided informed consent for biobank research, based on the Finnish Biobank Act. Alternatively, older research cohorts, collected prior the start of FinnGen (in August 2017), were collected based on study-specific consents and later transferred to the Finnish biobanks after approval by Fimea, the National Supervisory Authority for Welfare and Health. Recruitment protocols followed the biobank protocols approved by Fimea. The Coordinating Ethics Committee of the Hospital District of Helsinki and Uusimaa (HUS) approved the FinnGen study protocol Nr HUS/990/2017.

The FinnGen study is approved by Finnish Institute for Health and Welfare (THL), approval number THL/2031/6.02.00/2017, amendments THL/1101/5.05.00/2017, THL/341/6.02.00/2018, THL/2222/6.02.00/2018, THL/283/6.02.00/2019, THL/1721/5.05.00/2019, Digital and population data service agency VRK43431/2017-3, VRK/6909/2018-3, VRK/4415/2019-3 the Social Insurance Institution (KELA) KELA 58/522/2017, KELA 131/522/2018, KELA 70/522/2019, KELA 98/522/2019, and Statistics Finland TK-53-1041-17.

The Biobank Access Decisions for FinnGen samples and data utilized in FinnGen Data Freeze 5 include: THL Biobank BB2017_55, BB2017_111, BB2018_19, BB_2018_34, BB_2018_67, BB2018_71, BB2019_7, BB2019_8, BB2019_26, Finnish Red Cross Blood Service Biobank 7.12.2017, Helsinki Biobank HUS/359/2017, Auria Biobank AB17-5154, Biobank Borealis of Northern Finland_2017_1013, Biobank of Eastern Finland 1186/2018, Finnish Clinical Biobank Tampere MH0004, Central Finland Biobank 1-2017, and Terveystalo Biobank STB 2018001.

### Cohort pruning

The initial sample consisted of 541 PROFILE cases, 272 UK Biobank cases, and 302,081 UK Biobank controls (**Table S2**). We removed samples where there was a discordance between self-reported and X:Y coverage ratios, as well as samples with > 4% contamination according to VerifyBamID. The cohort was screened with KING to ensure that only unrelated (up to third-degree) individuals were included in the test. To reduce variation due to population stratification, we only included individuals with a probability of European Ancestry ≥ 0.98 based on PEDDY predictions and individuals within four standard deviations of principal components 1-4 (**Extended Data Fig. 1**). Further, samples were required to have greater than 95% of CCDS (release 22) bases covered with at least 10-fold coverage.

The control cohort was further restricted to include only individuals without a history of respiratory disease (**Table S1**). Using random sampling of the controls, we gender matched the control cohort to the case cohort (75% male). The final cohort consisted of 752 cases and 119,055 controls.

### Exome-wide association study (exWAS)

We tested 564,159 protein-coding variants association with IPF status. We specifically included all variants that were present in at least 12 individuals in the case-control cohort and passed the following QC criteria: minimum coverage 10X; percent of alternate reads in heterozygous variants ≥ 0.3 and ≤ 0.8; binomial test of alternate allele proportion departure from 50% in heterozygous state p > 10^−6^; genotype quality score (GQ) ≥ 30; Fisher’s strand bias score (FS) ≤ 200 (indels) ≤ 60 (SNVs); mapping quality score (MQ) ≥ 40; quality score (QUAL) ≥ 30; read position rank sum score (RPRS) ≥ −2; mapping quality rank sum score (MQRS) ≥ −8; DRAGEN variant status = PASS; Binomial test of difference in missingness between cases and controls p < 10^−6^; variant did not achieve Hardy-Weinberg Equilibrium Exact p < 10^−5^; variant site is not missing (i.e., <10X coverage) in ≥ 1% of cases or controls; variant did not fail above QC in ≥ 0.5% of cases or controls; variant site achieved 10-fold coverage in ≥ 50% of GnomAD exomes, and if variant was observed in GnomAD the variant calls in GnomAD achieved exome z-score ≥ −0.2 and exome MQ ≥ 30.

P-values were generated via Fisher’s exact test. The signal emerging from a novel *SPDL1* missense variant (p.Arg20Gln [rs116483731]) was confirmed in the FinnGen dataset (release 5), which includes 1,028 cases and 196,986 controls. We generated a combined p-value via Stouffer’s Z-test and defined genome-wide significance as the conventional p < 5 × 10^−8^.

### Collapsing analysis

To perform collapsing analyses, we aggregate variants within each gene that fit a given set of criteria, identified as “qualifying variants” (QVs)^6^. We performed 10 non-synonymous collapsing analyses, including 9 dominant and one recessive model, plus an additional synonymous variant model as a negative control. In each model, for each gene, the proportion of cases is compared to the proportion of controls carrying one or more qualifying variants in that gene. The exception is the recessive model, where a subject must have two qualifying alleles. The criteria for qualifying variants in each collapsing analysis model are in **Table S4**. P-values were generated via Fisher’s exact test.

For all models (**Table S4**) we applied the following QC filters: minimum coverage 10X; annotation in CCDS transcripts (release 22; ~34Mb); percent alternate reads in homozygous genotypes ≥ 0.8; percent of alternate reads in heterozygous variants ≥ 0.3 and ≤ 0.8; binomial test of alternate allele proportion departure from 50% in heterozygous state p > 10^−6^; genotype quality score (GQ) ≥ 30; Fisher’s strand bias score (FS) ≤ 200 (indels) ≤ 60 (SNVs); mapping quality score (MQ) ≥ 40; quality score (QUAL) ≥ 30; read position rank sum score (RPRS) ≥ −2; mapping quality rank sum score (MQRS) ≥ −8; DRAGEN variant status = PASS; Binomial test of difference in missingness between cases and controls p < 10^−6^; variant is not missing (i.e., <10X coverage) in ≥ 1% of cases or controls; variant site achieved 10-fold coverage in ≥ 25% of GnomAD samples, and if variant was observed in GnomAD the variant calls in GnomAD achieved exome z-score ≥ −2.0 and exome MQ ≥ 30.

### *SPDL1* haplotype analysis

We constructed haplotypes for the *SPDL1* locus in the PROFILE cohort (N=507) and in a random subset of 25,000 unrelated European individuals from the UK Biobank. This was done by phasing a total of 76 variants (MAF > 1%) that were located within a 10kb window of rs116483731. The genotype phasing was performed using MACH^18^, implementing the following parameters: --states 1000 and --rounds 60. Following the genotype phasing, we estimated the number of distinct haplotypes and their corresponding frequencies in each dataset. Next, in each dataset, we identified the *SPDL1* risk haplotypes i.e. haplotypes that contained the risk allele ‘A’ for rs116483731. Then, for each *SPDL1* risk haplotype that was identified, we determined the corresponding ancestral haplotype (having the reference allele ‘G’ in place of the risk allele ‘A’ for rs116483731) and estimated the percentage of those with that ancestral haplotype background who carried the rs116483731 mutation.

### Putatively Pathogenic *RTEL1*, *PARN*, *TERC* and *TERT* variants

To identify PROFILE subjects with putatively pathogenic variants in *RTEL1*, *PARN*, *TERT* or *TERC* we adopted the following criteria:

1. Variants were required to fulfil the following QC thresholds:
a. Percentage of variant allele reads ≥ 0.3
b. Binomial exact test of the departure from heterozygous expectation of 0.5 for variant allele read ratio p>0.001
c. GQ ≥ 30
d. QUAL ≥ 30
e. Variant affects a CCDS transcript
2. For putative protein-truncating variants (PTVs) in *RTEL1, PARN* and *TERT* the gnomAD minor allele frequency ≤ 0.05% (gnomAD popmax)
3. For missense variants in *TERT* and *PARN* the gnomAD minor allele frequency = 0 (ultra-rare)
4. For *TERC* noncoding RNA variants they were annotated by ClinVar as “Pathogenic” or the same *TERC* nucleotide was recurring affected by multiple variants.

All putatively pathogenic variants are included in **Table S9**.

### Effect size comparisons

We compared the effect sizes for different classes of variants implicated in IPF: PTVs in *TERT, RTEL1,* and *PARN*; putatively damaging missense variants in TERT, and GWAS loci that reached genome-wide significance in the largest IPF GWAS performed to date^4^. We used the UCSC Genome Browser LiftOver tool to convert the reported GRCh37 coordinates to GRCh38 coordinates, requiring 100% of bases to remap. We excluded the *MAPT* risk allele (rs2077551), as this allele failed the gnomAD random forest filter. For coding variants in *TERT, RTEL1,* and *PARN*, we used the case versus control odds ratios calculated in our collapsing analysis. For the GWAS loci, we used the PROFILE WGS data to calculate the frequency in cases, and we used the GnomAD non-Finnish European allele frequencies to derive the frequency in controls.

### Clinical characteristics

We compared clinical features for individuals in the PROFILE cohort who carried the *MUC5B* risk allele, the *SPDL1* risk allele, or putatively pathogenic variants in *RTEL1, TERC, TERT*, and *PARN* (**Table S6**). For each genotype, carriers were compared to all other individuals in the PROFILE cohort (i.e. non-carriers for the given risk allele). We specifically assessed differences in gender, survival months, sample age, height, weight, forced vital capacity (FVC), and diffusing capacity for carbon monoxide (DLCO). We performed median data-imputation to account for missing data. The p-values for comparing gender imbalances were generated via Fisher’s exact test, whereas p-values for all other clinical characteristics were generated via the Mann Whitney U test. Mortality rates were compared among carriers of rare variants in *TERT, TERC, PARN, and RTEL1,* carriers of the *SPDL1* risk allele, and carriers of the *MUC5B* risk allele versus non-carriers using the Kaplain-Meier method and p-values were generated using a log-rank test.

### Telomere length comparisons

For 96% of the samples the read lengths ranged between 148 to 150 bp. These BAM files were put through computational telomere length prediction method Telseq v0.0.1^12^ using a repeat number of 10. WGS sequences in this cohort did not use a PCR-free DNA sequencing protocol. Logistic regression was used to determine differences in telomere lengths between carriers of the *SPDL1* risk allele, carriers of the *MUC5B* risk allele, and carriers of rare variants in *TERT*, *TERC*, *RTEL1*, or *PARN*, adjusted for age and sex.

### Mantis-ml

Known pulmonary fibrosis-associated genes were automatically extracted from the Human Phenotype Ontology (HPO) by specifying the following term in the input configuration file of mantis-ml^8^: “Disease/Phenotype terms: pulmonary fibrosis”. This resulted in the following 38 HPO-defined seed genes: *ABCA3, SFTPA2, AP3B1, CAV1, DPP9, CFTR, CCR6, CCN2, CTLA4, PTPN22, TINF2, DKC1, DSP, FAM13A, DCTN4, RTEL1, HLA-DPA1, HLA-DPB1, HLA-DRB1, HPS1, IRF5, PARN, PRTN3, FAM111B, RCBTB1, STN1, SFTPA1, NOP10, CLCA4, SFTPC, NHP2, STX1A, ATP11A, MUC5B, TERC, TERT, TGFB1* and *HPS4*.

Automatic feature compilation on mantis-ml was performed by providing the following “Additional associated terms” in the input configuration file: “pulmonary, respirat and lung.” Mantis-ml was trained using six different classifiers: Extra Trees, XGBoost, Random Forest, Gradient Boosting, Support Vector Classifier and feed-forward Deep Neural Net.

Once the mantis-ml genome-wide probabilities of being an IPF gene were generated, we performed a hypergeometric test to determine whether the top-ranked collapsing analysis genes (i.e. genes achieving a p < 0.05 in the collapsing analyses) were significantly enriched for the top 5% of mantis-ml IPF-predicted genes. A statistically significant result from the hypergeometric test suggests that there are disease-ascertained genes among the top hits of the collapsing results. We then tested for this enrichment after excluding known IPF genes from our collapsing results to determine whether the signal was independent of genes already associated with IPF. In parallel, we also performed the hypergeometric enrichment test using the synonymous collapsing model to define our empirical null distribution. Among all six mantis-ml integrated classifiers used for training, Gradient Boosting achieved the highest enrichment signal against the collapsing analysis results from IPF.

## Acknowledgments

We thank the participants and investigators in the UK biobank study (Resource Application Number 26041), and PROFILE clinical colleagues who have referred IPF cases. We thank the AstraZeneca Centre for Genomics Research Analytics and Informatics team for processing and analysis of sequencing data. L.W. holds a GSK/British Lung Foundation Chair in Respiratory Research. The research was partially supported by the National Institute for Health Research (NIHR) Leicester Biomedical Research Centre; the views expressed are those of the author(s) and not necessarily those of the National Health Service (NHS), the NIHR or the Department of Health. P.L.M. is supported by an Action for Pulmonary Fibrosis Mike Bray fellowship. T.M.M. is supported by a National Institute for Health Research Clinician Scientist Fellowship (NIHR ref: CS-2013-13-017) and is a British Lung Foundation Chair in Respiratory Research (C17-3).

The FinnGen project is funded by two grants from Business Finland (HUS 4685/31/2016 and UH 4386/31/2016) and eleven industry partners (AbbVie Inc, AstraZeneca UK Ltd, Biogen MA Inc, Celgene Corporation, Celgene International II Sàrl, Genentech Inc, Merck Sharp & Dohme Corp, Pfizer Inc., GlaxoSmithKline, Sanofi, Maze Therapeutics Inc., Janssen Biotech Inc). Following biobanks are acknowledged for collecting the FinnGen project samples: Auria Biobank (www.auria.fi/biopankki), THL Biobank (www.thl.fi/biobank), Helsinki Biobank (www.helsinginbiopankki.fi), Biobank Borealis of Northern Finland (https://www.ppshp.fi/Tutkimus-ja-opetus/Biopankki/Pages/Biobank-Borealis-briefly-in-English.aspx), Finnish Clinical Biobank Tampere (www.tays.fi/en-US/Research_and_development/Finnish_Clinical_Biobank_Tampere), Biobank of Eastern Finland (www.ita-suomenbiopankki.fi/en), Central Finland Biobank (www.ksshp.fi/fi-FI/Potilaalle/Biopankki), Finnish Red Cross Blood Service Biobank (www.veripalvelu.fi/verenluovutus/biopankkitoiminta) and Terveystalo Biobank (www.terveystalo.com/fi/Yritystietoa/Terveystalo-Biopankki/Biopankki/). All Finnish Biobanks are members of BBMRI.fi infrastructure (www.bbmri.fi).

