## Supplementary Material for "Identification of a novel missense variant in *SPDL1* associated with idiopathic pulmonary fibrosis"

#### Extended data

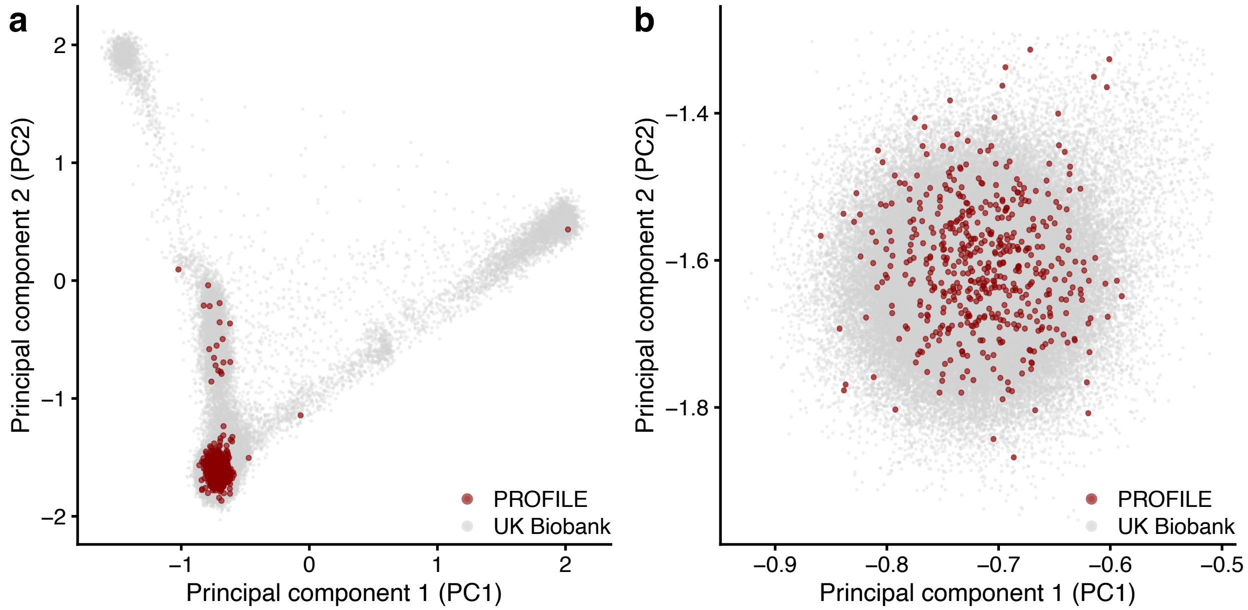

**Supplementary Figure 1. Principal component plot of ancestry axes. (a)** Principal components (PCs) 1 and 2 for 133,056 samples contained in the UK Biobank and 530 samples from the PROFILE cohort. **(b)** Samples retained after filtering for individuals with Peddy-based probability of being European was greater than 0.98 and the sample was within the quadrant boundary defined by: [PC1: -0.9280 to -0.5092] and [PC2: -1.953 to -1.288].

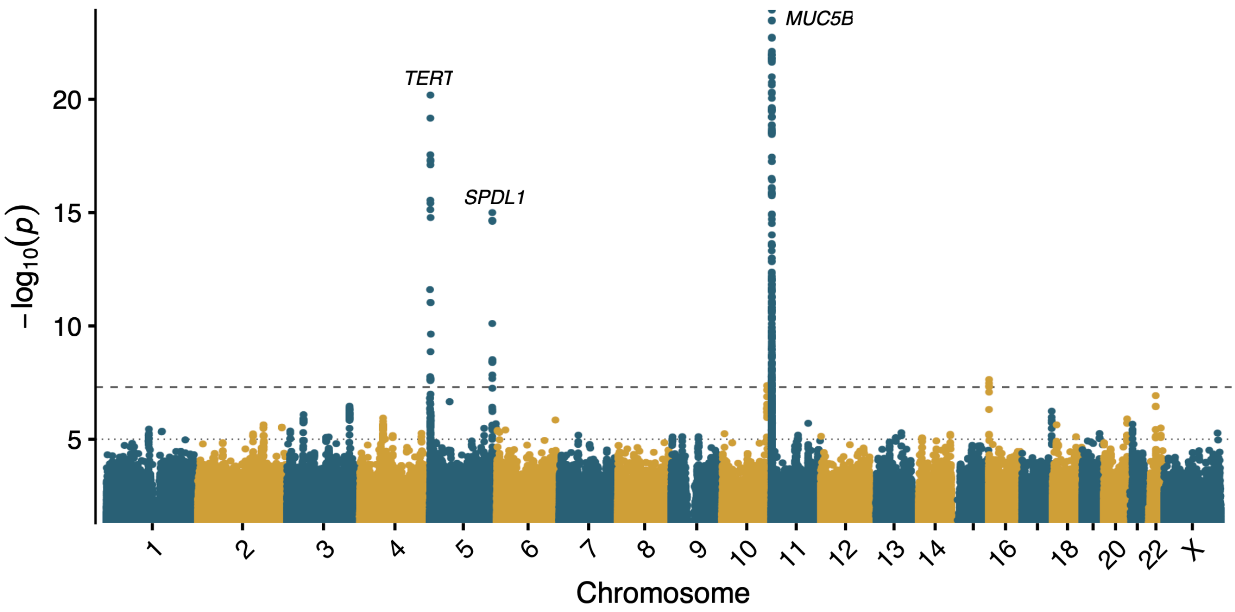

**Supplementary Figure 2. Association of single-nucleotide variants with IPF in the FinnGen cohort.** Manhattan plot depicting *p*-values of the 16,380,412 tested for association with IPF status in the FinnGen cohort (release 5). The y-axis has been capped at 25.

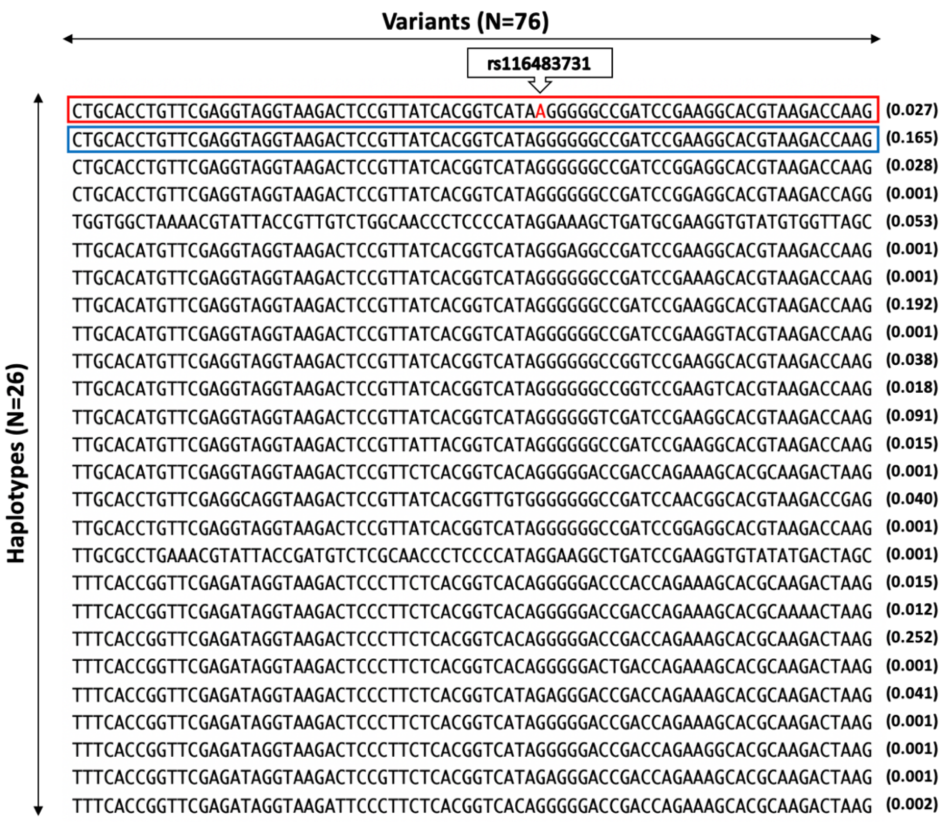

**Supplementary Figure 3. Haplotype analysis**. Depiction of the 26 distinct haplotypes that were identified for 76 variants within a 10 kilobase window of the *SPDL1* index variant rs116483731 in the PROFILE cohort. The risk allele ‘A’ for rs116483731 was present on only one of the 26 haplotypes (red box), while the remaining haplotypes all harboured the ancestral “G” allele at this position. The blue box represents the corresponding ancestral haplotype, which was present in 16.5% of the cohort. The numbers in parentheses adjacent to each haplotype represents the observed frequency in the cohort.

We further confirmed the finding of a shared common haplotype for the rs116483731 variant using an independent population-based dataset of 25,000 individuals sampled from the UK Biobank and found >99% of the rs116483731 risk alleles had an identical ancestral haplotype structure to the *SPDL1* risk haplotype observed among Imperial IPF cases.

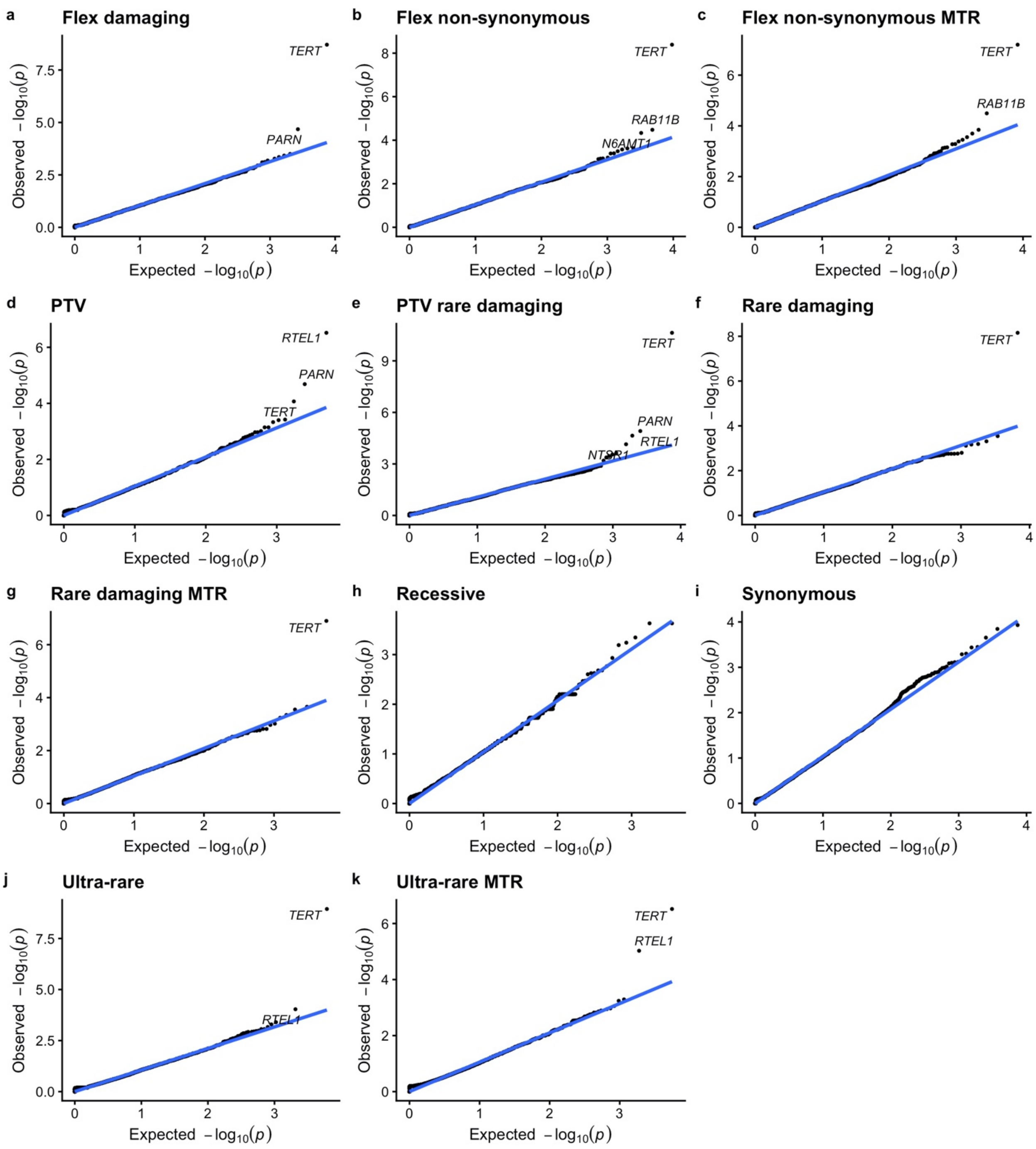

**Supplementary Figure 4**. **Gene-level collapsing results. (a-k)** Quantile-quantile plots for each gene-level collapsing model. Each model included 18,665 protein-coding genes. No novel genes achieved study-wide significance (adjusted $\alpha$ < .05 / [18,665 x 10] = 2.7 x $10^{-7}$).

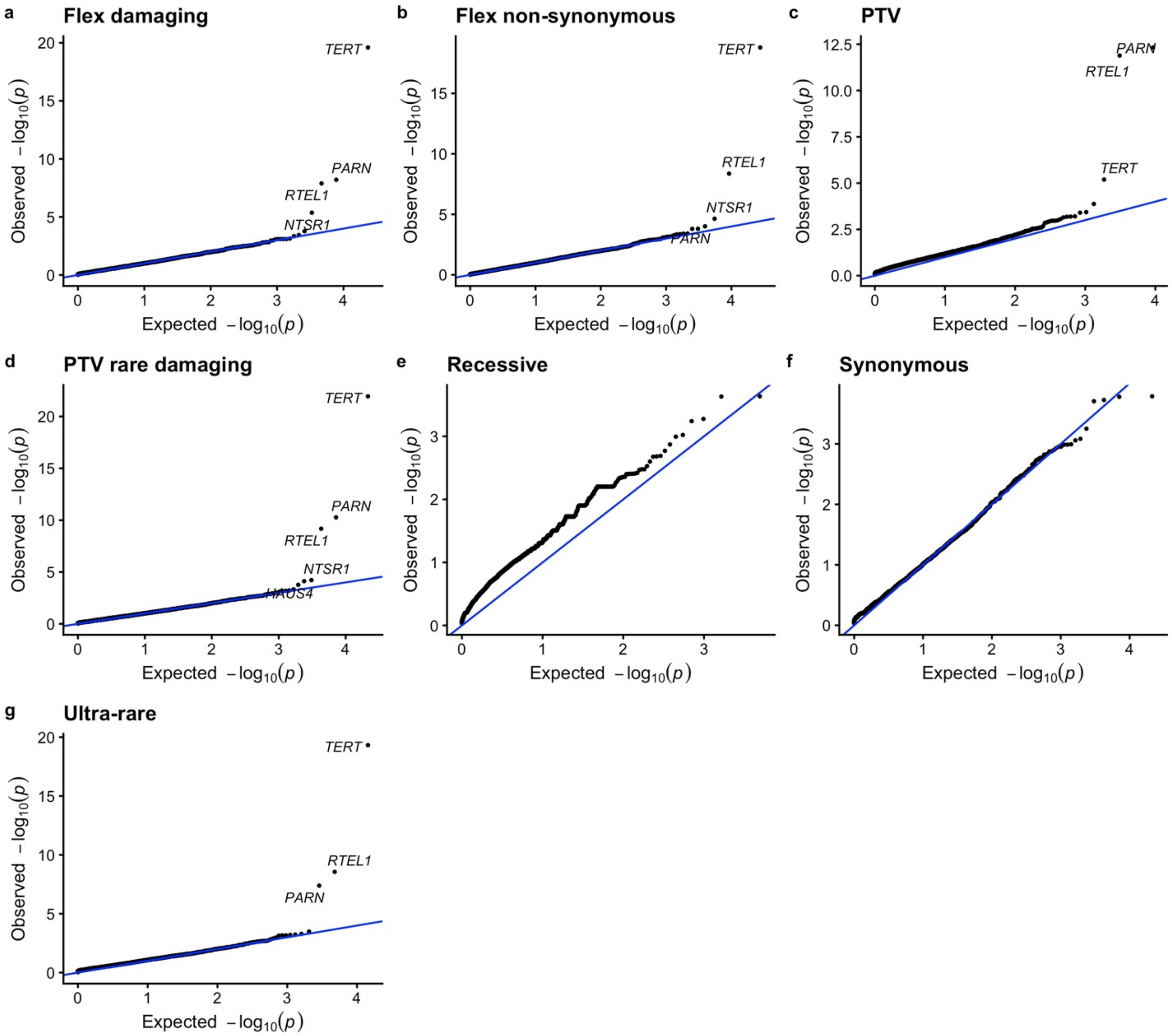

**Supplementary Figure 5**. **Combined analysis for gene-level collapsing results. (a-g)** Quantile-quantile plots for each gene-level collapsing model, in which we combined data from the present study with a previous IPF collapsing study[4]. Each model included 18,665 protein-coding genes. No novel genes achieved significance.

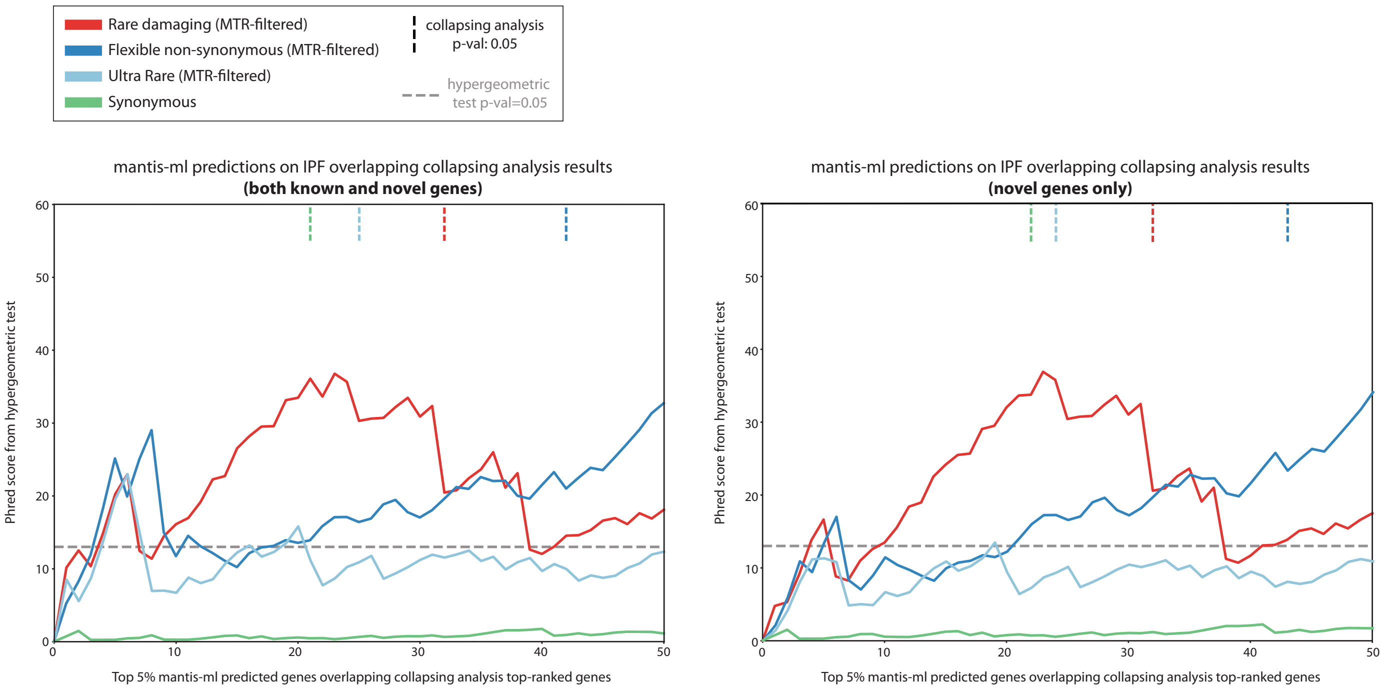

| | Min. p-value (up to last significant gene) | | p-value at last significant gene from collapsing | |
|---|---|---|---|---|
| **QV model** | All genes | Novel genes only | All genes | Novel genes only |
| Rare damaging | **0.00021** | **0.000203** | **0.00899** | **0.00873** |
| Flex non-synonymous | **0.00124** | **0.00267** | **0.00792** | **0.00468** |
| Ultra-rare damaging | **0.00504** | **0.0454** | 0.0814 | 0.118 |
| Synonymous | 0.715 | 0.713 | 0.902 | 0.856 |

**Supplementary Figure 6. Cross-validation of *mantis-ml* predictions with cohort-level rare-variant association studies.** Hypergeometric test enrichment of IPF-specific *mantis-ml* predictions against three collapsing models: “rare damaging MTR,” “flexible non-synonymous MTR,” and “ultra-rare damaging MTR.” The synonymous collapsing model is included as a negative control. The horizontal dashed grey line corresponds to the significance threshold of p = 0.05 for the hypergeometric tests, with signals above this line indicating a significant enrichment. The vertical dashed lines indicate the p-value of the top-ranked gene from each collapsing model achieving a p-value < 0.05. The plot on the right depicts the same analysis except that known IPF genes were removed. The table below includes the hypergeometric test p-values (considering either all genes or novel genes only).

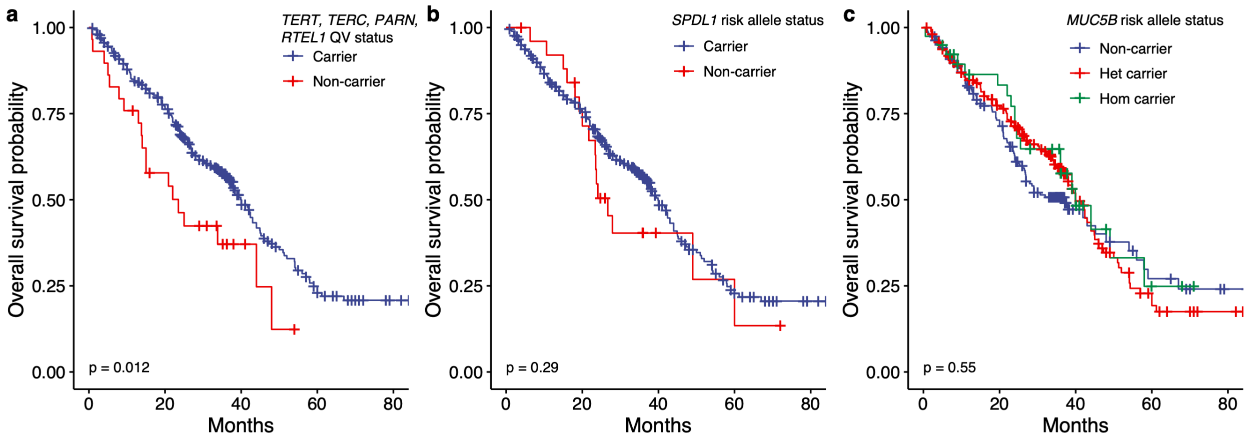

**Supplementary Figure 7. Kaplan-Meier survival curves of PROFILE cases stratified by genetic risk factor. (a)** Comparison of cases with qualifying rare variants (QVs) in *TERT*, *TERC, PARN,* and *RTEL1* versus non-carriers. **(b)** Comparison of cases carrying the *SPDL*1 risk allele versus non-carriers. **(c)** Comparison of *MUC5B* risk allele carriers versus the remainder of the PROFILE cohort. Log-rank p-values are indicated on each plot.

| Exclusion Criteria | UK Biobank Field |
|---|---|
| ICD10 Chapter X Diseases of the respiratory system (root node ID 10) | 41270 |
| ICD10 Chapter XVIII Symptoms, signs and abnormal clinical and laboratory findings, not elsewhere classified (root node 18; Respiratory Block R05-R06, R08-R09) | 41270 |
| ICD9 Chapter VIII Diseases of the respiratory system (460-519) | 41271 |
| ICD10 Chapter X Diseases of the respiratory system (root node ID 10) | 40001 |
| keywords – “pulmon,” “respire,” “asthma,” “airways,” “bronchi,” “interstitial lung,” “pneumonia,” “asbestosis,” “pneumothorax,” “lung abscess,” “emphysema,” “empyema,” “pleural plaques,” “pleural effusion,” “alveolitis” | 40010 |
| Self-declared respiratory (root node ID 1072) | 20002 |
| Online follow-up > Work environment > Medical information > Doctor Diagnosed | 22127 - 22141 |

**Supplementary Table 1**: Exclusion criteria used for screening UK Biobank controls.

| Inclusion Criteria | PROFILE | UK Biobank IPF | UK Biobank Controls |
|---|---|---|---|
| Initial Cohort | 541 (100%) | 272 (100%) | 302,081 |
| Restrict controls to non-respiratory disease (Table S2) | 541 (100%) | 272 (100%) | 219,704 (72.7%) |
| Contamination FREEMIX < 4% based on VerifyBamID | 540 (99.8%) | 270 (99.3%) | 219,627 (72.7%) |
| Gender concordance between reported and genetic gender | 534 (98.7%) | 270 (99.3%) | 219,627 (72.7%) |
| ≥95% of CCDS r22 bases covered with ≥10-fold coverage | 534 (98.7%) | 270 (99.3%) | 219,622 (72.7%) |
| Probability of European ancestry ≥ 0.98 based on PEDDY | 512 (94.6%) | 251 (92.3%) | 206,938 (68.5%) |
| Within 4 standard deviations (SD) of PC 1-4 for probability of European ancestry mean | 511 94.5% | 251 (92.3%) | 206,415 (68.2%) |
| Cryptic Relatedness Pruning (up to 3rd degree based on KING) | 507 (93.7%) | 245 (90.1%) | 200,203 (66.3%) |
| Within ±4SD of Novel CCDS SNV mean | 507 (93.7%) | 245 (90.1%) | 199,963 (66.2%) |
| Gender match controls (Male 75%) | 507 (93.7%) | 245 (90.1%) | 119,055 (39.4%) |
| **Final Test Cohort** | **507** (93.7%) | **245** (90.1%) | **119,055** (39.4%) |

**Supplementary Table 2**: Sample-level quality control filtering and case-control harmonization.

| | UK | | | | | | Colorado | | | | Chicago | | | | Meta | |
|---|---|---|---|---|---|---|---|---|---|---|---|---|---|---|---|---|
| | **Ref allele** | **Effect allele** | **MAF** | **Rsq** | **OR [95% CI]** | **p** | **MAF** | **Rsq** | **OR [95% CI]** | **p** | **MAF** | **Rsq** | **OR [95% CI]** | **p** | **OR [95% CI]** | **p** |
| rs116483731 | G | A | 0.8% | 0.84 | 1.38 [0.70, 2.74] | 0.354 | 0.9% | 0.81 | 3.95 [2.49, 6.28] | $6.01 \times 10^{-9}$ | 1.2% | 0.83 | 1.19 [0.55, 2.61] | 0.656 | 2.40 [1.70, 3.40] | $7.55 \times 10^{-7}$ |

**Supplementary Table 3**: P-values for the *SPDL1* risk variant (rs116483731) from the latest GWAS meta-analysis of IPF risk alleles, which included some overlapping samples from the PROFILE study[4]. This variant did not achieve genome-wide significance in this GWAS but demonstrated a similar effect size. Allen et al. required P < 0.05 in all 3 contributing studies to be tested in the replication cohort.

| Collapsing model | GnomAD MAF | Internal MAF | Variant type | REVEL[3] cutoff | MTR cutoff |
|---|---|---|---|---|---|
| **Synonymous (negative control)** | ≤ 0.005% | ≤ 0.05% | Synonymous | - | - |
| **PTV (Protein Truncating Variant)** | ≤ 0.1% (popmax) | ≤ 0.1% | PTV | - | - |
| **Ultra-rare damaging** | <0.0004% | ≤ 0.025% | PTV, missense, inframe indels | ≥ 0.25 | - |
| **Ultra-rare damaging MTR** | <0.0004% | ≤ 0.025% | PTV, missense, inframe indels | ≥ 0.25 | MTR ≤ 25th percentile or Intragenic MTR ≤ 50th %ile |
| **Rare damaging** | ≤ 0.005% | ≤ 0.05% | missense | ≥ 0.25 | - |
| **Rare damaging MTR** | ≤ 0.005% | ≤ 0.05% | missense | ≥ 0.25 | MTR ≤ 25th percentile or Intragenic MTR ≤ 50th %ile |
| **Flexible damaging** | ≤ 0.05% (global)<br>≤ 0.1% (popmax) | ≤ 0.1% | PTV, missense, inframe indels | ≥ 0.25 | - |
| **Flexible non-synonymous** | ≤ 0.05% (global)<br>≤ 0.1% (popmax) | ≤ 0.1% | PTV, missense, inframe indels | - | - |
| **Flexible non-synonymous MTR** | ≤ 0.05% (global)<br>≤ 0.1% (popmax) | ≤ 0.1% | PTV, missense, inframe indels | - | MTR ≤ 25th percentile or Intragenic MTR ≤ 50th %ile |
| **PTV or rare damaging missense** | PTV ≤ 0.1%<br>missense ≤ 0.005% (global)<br>missense ≤ 0.05% (popmax) | PTV ≤ 0.1%<br>missense ≤ 0.05% | PTV, missense, inframe indels | ≥ 0.25 | - |
| **Recessive non-synonymous** | ≤ 0.5% (popmax) | ≤ 0.5% | PTV, missense, inframe indels | - | - |

**Supplementary Table 4.** Collapsing models. MTR = missense tolerance ratio[2]. REVEL and MTR cutoffs only apply to missense variants.

< Table_S5_collapsing_supplement.xlsx>

**Supplementary Table 5.** Results from 11 different collapsing models.

< Table_S6_all_cmh_results.xlsx>

**Supplementary Table 6.** Results from combined collapsing analyses for 6 different models.

| Field | All | *MUC5B* (rs35705950) | p-value (vs. others) | *SPDL1* (p.Arg20Gln) | p-value (vs. others) | *TERT, TERC, RTEL1, PARN* (rare variant) | p-value (vs. others) |
|---|---|---|---|---|---|---|---|
| Sample size | 507 | 309 | N/A | 26 | N/A | 31 | N/A |
| Male gender | 387 | 232 | 0.454 | 17 | 0.233 | 17 | 0.007 |
| Survival months* (median) | 21.1 | 22.5 | 0.046 | 22.55 | 0.569 | 14 | 0.067 |
| Sample age (median) | 71 | 71 | 0.127 | 69.5 | 0.379 | **66** | **0.0008** |
| Height (median) | 171 | 171 | 0.212 | 171 | 0.348 | 169.5 | 0.103 |
| Weight (median) | 82.6 | 82.6 | 0.159 | 79.25 | 0.287 | 82.6 | 0.968 |
| FVC pp (median) | 76.86 | 76.79 | 0.16 | 77.94 | 0.49 | 76.82 | 0.35 |
| DLCO pp (median) | 47.9 | 47.68 | 0.088 | 46.52 | 0.981 | 47.9 | 0.958 |
| TelSeq (median) | 0.77 | 0.77 | 0.180 | 0.77 | 0.947 | **0.68** | **0.002** |
| Family history | 42 | 26 | 1 | 4 | 0.259 | 3 | 0.734 |

**Supplementary Table 7. Clinical characteristics of PROFILE cohort.** P-values for gender were generated via Fisher's exact test. P-values for values with reported medians were calculated using the Mann-Whitney U test. Bonferroni corrected p-values ≤ 0.003 are bolded. *Survival months were only considered for the 238 samples with reported mortality data (deceased).

| **Covariate** | ***MUC5B*** (rs35705950) | ***SPDL1*** (p.Arg20gln) | ***TERT, TERC, RTEL1, PARN*** (rare variant) |
|---|---|---|---|
| Age Odds Ratio (p-value) | 1 (0.03) | 0.98 (0.4) | 0.94 (0.002) |
| Gender Odds Ratio (p-value) | 0.84 (0.4) | 0.58 (0.2) | 0.32 (0.004) |
| TelSeq-inferred telomere length (kb) Odds Ratio (p-value) | 0.86 (0.2) | 0.62 (0.6) | 0.03 (0.004) |

**Supplementary Table 8. TelSeq logistic regression analysis.** Odds ratios with p-values in parentheses

| Sample Name | Variant ID | Function | Gene Name | Transcript codon change | Transcript AA change | gnomAD Exome Alleles | ClinVar Disease | ClinVar Clinical Signif. | Family history |
|---|---|---|---|---|---|---|---|---|---|
| PROFILE1 | 16-14482828-C-T | splice acceptor | *PARN* | c.1298-1G>A | NA | 1 | NA | NA | Yes |
| PROFILE2 | 16-14555652-A-G | splice donor | *PARN* | c.1135+2T>C | NA | NA | NA | NA | |
| PROFILE3 | 16-14617651-C-T | splice acceptor | *PARN* | c.145-1G>A | NA | NA | NA | NA | |
| PROFILE4 | 16-14629675-C-T | splice acceptor | *PARN* | c.-164-1G>A | NA | NA | NA | NA | |
| PROFILE5 | 16-14447001-CCT-C | frameshift | *PARN* | c.1566 1567delAG | p.Glu524fs | NA | NA | NA | |
| PROFILE6 | 16-14482683-TG-T | frameshift | *PARN* | c.1441delC | p.Gln481fs | 1 | NA | NA | |
| PROFILE7 | 16-14482683-TG-T | frameshift | *PARN* | c.1441delC | p.Gln481fs | 1 | NA | NA | |
| PROFILE8 | 16-14628170-GC-G | splice donor | *PARN* | c.-7+1delG | NA | NA | NA | NA | |
| PROFILE9 | 16-14555694-C-A | missense | *PARN* | c.1095G>T | p.Arg365Ser | NA | NA | NA | |
| PROFILE10 | 16-14593338-A-G | missense | *PARN* | c.698T>C | p.Met233Thr | NA | NA | NA | |
| PROFILE11 | 16-14627163-G-C | missense | *PARN* | c.87C>G | p.Phe29Leu | NA | NA | NA | |
| PROFILE12 | 20-63690172-C-T | stop gained | *RTEL1* | c.1558C>T | p.Arg520* | NA | NA | NA | |
| PROFILE13 | 20-63693247-C-T | stop gained | *RTEL1* | c.2287C>T | p.Arg763* | 20 | Dyskeratosis congenita | P/LP | |
| PROFILE14 | 20-63693247-C-T | stop gained | *RTEL1* | c.2287C>T | p.Arg763* | 20 | Dyskeratosis congenita | P/LP | |
| PROFILE15 | 20-63679873-AG-A | frameshift | *RTEL1* | c.394delG | p.Ala132fs | NA | NA | NA | |
| PROFILE16 | 20-63689531-GA-G | frameshift | *RTEL1* | c.1241delA | p.Asn414fs | NA | NA | NA | |
| PROFILE17 | 20-63694447-ACCAGGGCAGGCCCCACCTGTCGC-A | frameshift | *RTEL1* | c.2405 2427delGCAGGCCCCACCTGTCGCCCAGG | p.Gly802fs | NA | NA | NA | Yes |
| PROFILE18 | 20-63693211-C-T | stop gained | *RTEL1* | c.2251C>T | p.Arg751* | 7 | Dyskeratosis congenita \| Idiopathic fibrosing alveolitis \| Pulmonary fibrosis and/or bone marrow failure | P/LP | |
| PROFILE19 | 5-1260499-C-T | missense | *TERT* | c.2945G>A | p.Cys982Tyr | NA | NA | NA | |
| PROFILE20 | 5-1268521-C-T | missense | *TERT* | c.2581G>A | p.Gly861Arg | NA | Idiopathic fibrosing alveolitis \| Dyskeratosis congenita | VUS | |
| PROFILE21 | 5-1268577-C-T | missense | *TERT* | c.2525G>A | p.Cys842Tyr | NA | NA | NA | |

| | | | | | | | | | |
|---|---|---|---|---|---|---|---|---|---|
| PROFILE22 | 5-1268577-C-T | missense | *TERT* | c.2525G>A | p.Cys842Tyr | NA | NA | NA | |
| PROFILE23 | 5-1271156-G-A | missense | *TERT* | c.2431C>T | p.Arg811Cys | 0 | Dyskeratosis congenita | P/LP | |
| PROFILE24 | 5-1272184-C-G | splice donor | *TERT* | c.2382+1G>C | NA | NA | NA | NA | |
| PROFILE25 | 5-1278655-C-G | missense | *TERT* | c.2272G>C | p.Ala758Pro | NA | NA | NA | |
| PROFILE26 | 5-1278688-C-T | missense | *TERT* | c.2239G>A | p.Val747Ile | NA | Idiopathic fibrosing alveolitis \| Dyskeratosis congenita | VUS | |
| PROFILE27 | 5-1280244-G-A | missense | *TERT* | c.1864C>T | p.Arg622Cys | NA | NA | NA | |
| PROFILE28 | 5-1282573-G-A | missense | *TERT* | c.1625C>T | p.Ala542Val | NA | NA | NA | Yes |
| PROFILE29 | 5-1294213-C-T | missense | *TERT* | c.673G>A | p.Gly225Arg | NA | NA | NA | |
| PROFILE30 | 5-1293522-T-A | missense | *TERT* | c.1364A>T | p.His455Leu | 0 | NA | NA | |
| UKB 1 | 5-1254482-C-T | missense | *TERT* | c.3181G>A | p.Ala1061Thr | NA | NA | NA | |
| UKB 2 | 5-1279341-C-T | missense | *TERT* | c.2080G>A | p.Val694Met | 0 | Aplastic anemia \| Pulmonary fibrosis and/or bone marrow failure | P/LP | |
| UKB 3 | 5-1282573-G-A | missense | *TERT* | c.1625C>T | p.Ala542Val | NA | NA | NA | |
| UKB 4 | 5-1294277-C-G | missense | *TERT* | c.609G>C | p.Trp203Cys | NA | NA | NA | |
| UKB 5 | 5-1293499-CG-C | frameshift | *TERT* | c.1386delC | p.Tyr462fs | NA | NA | NA | |
| UKB 6 | 5-1294226-ACC-A | frameshift | *TERT* | c.658 659delGG | p.Gly220fs | 1 | NA | NA | |
| UKB 7 | 5-1294951-CAGGG-C | frameshift | *TERT* | c.35 38delCCCT | p.Ser12fs | NA | NA | NA | |
| UKB 8 | 16-14617596-G-A | stop gained | *PARN* | c.199C>T | p.Arg67* | 1 | NA | NA | |
| UKB 9 | 16-14617596-G-A | stop gained | *PARN* | c.199C>T | p.Arg67* | 1 | NA | NA | |
| UKB 10 | 16-14617596-G-A | stop gained | *PARN* | c.199C>T | p.Arg67* | 1 | NA | NA | |
| UKB 11 | 20-63694419-C-T | stop gained | *RTEL1* | c.2371C>T | p.Gln791* | NA | NA | NA | |
| UKB 12 | 20-63695374-GTCCT-G | frameshift | *RTEL1* | c.2883 2886delCCTT | p.Phe961fs | NA | NA | NA | |
| UKB 13 | 16-14593324-G-C | missense | *PARN* | c.712C>G | p.Gln238Glu | NA | NA | NA | |
| UKB 14 | 16-14617590-C-G | missense | *PARN* | c.205G>C | p.Gly69Arg | NA | NA | NA | |
| PROFILE31 | 3-169765024-T-C | non coding transcript exon variant | *TERC* | n.24A>G | NA | NA | Dyskeratosis congenita | VUS | |
| PROFILE32 | 3-169765024-T-A | non coding transcript exon variant | *TERC* | n.24A>T | NA | NA | NA | NA | |

| | | | | | | | | |
|---|---|---|---|---|---|---|---|---|
| PROFILE33 | 3-169765026-G-A | non coding transcript exon variant | *TERC* | n.22C>T | NA | NA | Dyskeratosis congenita | P/LP |

**Supplementary Table 9. Putatively Pathogenic *RTEL1*, *PARN*, *TERC* and *TERT* variants**

### Contributors of FinnGen

#### Steering Committee

Aarno Palotie Institute for Molecular Medicine Finland, HiLIFE, University of Helsinki, Finland
Mark Daly Institute for Molecular Medicine Finland, HiLIFE, University of Helsinki, Finland

**Pharmaceutical companies**
Howard Jacob Abbvie, Chicago, IL, United States
Athena Matakidou Astra Zeneca, Cambridge, United Kingdom
Heiko Runz Biogen, Cambridge, MA, United States
Sally John Biogen, Cambridge, MA, United States
Robert Plenge Celgene, Summit, NJ, United States
Mark McCarthy Genentech, San Francisco, CA, United States
Julie Hunkapiller Genentech, San Francisco, CA, United States
Meg Ehm GlaxoSmithKline, Brentford, United Kingdom
Dawn Waterworth GlaxoSmithKline, Brentford, United Kingdom
Caroline Fox Merck, Kenilworth, NJ, United States
Anders Malarstig Pfizer, New York, NY, United States
Kathy Klinger Sanofi, Paris, France
Kathy Call Sanofi, Paris, France
Tim Behrens Maze Therapeutics, San Francisco, CA, United States
Patrick Loerch Janssen Biotech, Beerse, Belgium

**University of Helsinki & Biobanks**
Tomi Mäkelä HiLIFE, University of Helsinki, Finland, Finland
Jaakko Kaprio Institute for Molecular Medicine Finland, HiLIFE, Helsinki, Finland, Finland
Petri Virolainen Auria Biobank / University of Turku / Hospital District of Southwest Finland, Turku, Finland
Kari Pulkki Auria Biobank / University of Turku / Hospital District of Southwest Finland, Turku, Finland
Terhi Kilpi THL Biobank / The National Institute of Health and Welfare Helsinki, Finland
Markus Perola THL Biobank / The National Institute of Health and Welfare Helsinki, Finland
Jukka Partanen Finnish Red Cross Blood Service / Finnish Hematology Registry and Clinical Biobank, Helsinki, Finland
Anne Pitkäranta Helsinki Biobank / Helsinki University and Hospital District of Helsinki and Uusimaa, Helsinki
Riitta Kaarteenaho Northern Finland Biobank Borealis / University of Oulu / Northern Ostrobothnia Hospital District, Oulu, Finland
Seppo Vainio Northern Finland Biobank Borealis / University of Oulu / Northern Ostrobothnia Hospital District, Oulu, Finland
Miia Turpeinen Northern Finland Biobank Borealis / University of Oulu / Northern Ostrobothnia Hospital
Raisa Serpi Northern Finland Biobank Borealis / University of Oulu / Northern Ostrobothnia Hospital
Tarja Laitinen Finnish Clinical Biobank Tampere / University of Tampere / Pirkanmaa Hospital District, Tampere, Finland
Johanna Mäkelä Finnish Clinical Biobank Tampere / University of Tampere / Pirkanmaa Hospital District, Tampere, Finland
Veli-Matti Kosma Biobank of Eastern Finland / University of Eastern Finland / Northern Savo Hospital District, Kuopio, Finland
Urho Kujala Central Finland Biobank / University of Jyväskylä / Central Finland Health Care District, Jyväskylä, Finland

**Other Experts/ Non-Voting Members**

| | |
|---|---|
| Outi Tuovila | Business Finland, Helsinki, Finland |
| Minna Hendolin | Business Finland, Helsinki, Finland |
| Raimo Pakkanen | Business Finland, Helsinki, Finland |

#### Scientific Committee

**Pharmaceutical companies**

| | |
|---|---|
| Jeff Waring | Abbvie, Chicago, IL, United States |
| Bridget Riley-Gillis | Abbvie, Chicago, IL, United States |
| Athena Matakidou | Astra Zeneca, Cambridge, United Kingdom |
| Heiko Runz | Biogen, Cambridge, MA, United States |
| Jimmy Liu | Biogen, Cambridge, MA, United States |
| Shameek Biswas | Celgene, Summit, NJ, United States |
| Julie Hunkapiller | Genentech, San Francisco, CA, United States |
| Dawn Waterworth | GlaxoSmithKline, Brentford, United Kingdom |
| Meg Ehm | GlaxoSmithKline, Brentford, United Kingdom |
| Dorothee Diogo | Merck, Kenilworth, NJ, United States |
| Caroline Fox | Merck, Kenilworth, NJ, United States |
| Anders Malarstig | Pfizer, New York, NY, United States |
| Catherine Marshall | Pfizer, New York, NY, United States |
| Xinli Hu | Pfizer, New York, NY, United States |
| Kathy Call | Sanofi, Paris, France |
| Kathy Klinger | Sanofi, Paris, France |
| Matthias Gossel | Sanofi, Paris, France |
| Robert Graham | Maze Therapeutics, San Francisco, CA, United States |
| Tim Behrens | Maze Therapeutics, San Francisco, CA, United States |
| Beryl Cummings | Maze Therapeutics, San Francisco, CA, United States |
| Wilco Fleuren | Janssen Biotech, Beerse, Belgium |

**University of Helsinki & Biobanks**

| | |
|---|---|
| Samuli Ripatti | Institute for Molecular Medicine Finland, HiLIFE, Helsinki, Finland |
| Johanna Schleutker | Auria Biobank / Univ. of Turku / Hospital District of Southwest Finland, Turku, Finland |
| Markus Perola | THL Biobank / The National Institute of Health and Welfare Helsinki, Finland |
| Mikko Arvas | Finnish Red Cross Blood Service / Finnish Hematology Registry and Clinical Biobank, Helsinki, Finland |
| Olli Carpén | Helsinki Biobank / Helsinki University and Hospital District of Helsinki and Uusimaa, Helsinki |
| Reetta Hinttala | Northern Finland Biobank Borealis / University of Oulu / Northern Ostrobothnia Hospital District, Oulu, Finland |
| Johannes Kettunen | Northern Finland Biobank Borealis / University of Oulu / Northern Ostrobothnia Hospital District, Oulu, Finland |
| Johanna Mäkelä | Finnish Clinical Biobank Tampere / University of Tampere / Pirkanmaa Hospital District, Tampere, Finland |
| Arto Mannermaa | Biobank of Eastern Finland / University of Eastern Finland / Northern Savo Hospital District, Kuopio, Finland |
| Jari Laukkanen | Central Finland Biobank / University of Jyväskylä / Central Finland Health Care District, Jyväskylä, Finland |
| Urho Kujala | Central Finland Biobank / University of Jyväskylä / Central Finland Health Care District, Jyväskylä, Finland |

**Other Experts/ Non-Voting Members**

| | |
|---|---|
| Outi Tuovila | Business Finland, Helsinki, Finland |
| Minna Hendolin | Business Finland, Helsinki, Finland |
| Raimo Pakkanen | Business Finland, Helsinki, Finland |

#### Clinical Groups

##### Neurology Group

| | |
|---|---|
| Hilkka Soininen | Northern Savo Hospital District, Kuopio, Finland |
| Valtteri Julkunen | Northern Savo Hospital District, Kuopio, Finland |
| Anne Remes | Northern Ostrobothnia Hospital District, Oulu, Finland |
| Reetta Kälviäinen | Northern Savo Hospital District, Kuopio, Finland |
| Mikko Hiltunen | Northern Savo Hospital District, Kuopio, Finland |
| Jukka Peltola | Pirkanmaa Hospital District, Tampere, Finland |
| Pentti Tienari | Hospital District of Helsinki and Uusimaa, Helsinki, Finland |
| Juha Rinne | Hospital District of Southwest Finland, Turku, Finland |
| Adam Ziemann | Abbvie, Chicago, IL, United States |
| Jeffrey Waring | Abbvie, Chicago, IL, United States |
| Sahar Esmaeeli | Abbvie, Chicago, IL, United States |
| Nizar Smaoui | Abbvie, Chicago, IL, United States |
| Anne Lehtonen | Abbvie, Chicago, IL, United States |
| Susan Eaton | Biogen, Cambridge, MA, United States |
| Heiko Runz | Biogen, Cambridge, MA, United States |
| Sanni Lahdenperä | Biogen, Cambridge, MA, United States |
| Janet van Adelsberg | Celgene, Summit, NJ, United States |
| Shameek Biswas | Celgene, Summit, NJ, United States |
| John Michon | Genentech, San Francisco, CA, United States |
| Geoff Kerchner | Genentech, San Francisco, CA, United States |
| Julie Hunkapiller | Genentech, San Francisco, CA, United States |
| Natalie Bowers | Genentech, San Francisco, CA, United States |
| Edmond Teng | Genentech, San Francisco, CA, United States |
| John Eicher | Merck, Kenilworth, NJ, United States |
| Vinay Mehta | Merck, Kenilworth, NJ, United States |
| Padhraig Gormley | Merck, Kenilworth, NJ, United States |
| Kari Linden | Pfizer, New York, NY, United States |
| Christopher Whelan | Pfizer, New York, NY, United States |
| Fanli Xu | GlaxoSmithKline, Brentford, United Kingdom |
| David Pulford | GlaxoSmithKline, Brentford, United Kingdom |

##### Gastroenterology Group

| | |
|---|---|
| Martti Färkkilä | Hospital District of Helsinki and Uusimaa, Helsinki, Finland |
| Sampsa Pikkarainen | Hospital District of Helsinki and Uusimaa, Helsinki, Finland |
| Airi Jussila | Pirkanmaa Hospital District, Tampere, Finland |
| Timo Blomster | Northern Ostrobothnia Hospital District, Oulu, Finland |
| Mikko Kiviniemi | Northern Savo Hospital District, Kuopio, Finland |
| Markku Voutilainen | Hospital District of Southwest Finland, Turku, Finland |
| Bob Georgantas | Abbvie, Chicago, IL, United States |
| Graham Heap | Abbvie, Chicago, IL, United States |
| Jeffrey Waring | Abbvie, Chicago, IL, United States |
| Nizar Smaoui | Abbvie, Chicago, IL, United States |
| Fedik Rahimov | Abbvie, Chicago, IL, United States |
| Anne Lehtonen | Abbvie, Chicago, IL, United States |
| Keith Usiskin | Celgene, Summit, NJ, United States |
| Tim Lu | Genentech, San Francisco, CA, United States |
| Natalie Bowers | Genentech, San Francisco, CA, United States |
| Danny Oh | Genentech, San Francisco, CA, United States |
| John Michon | Genentech, San Francisco, CA, United States |
| Vinay Mehta | Merck, Kenilworth, NJ, United States |
| Kirsi Kalpala | Pfizer, New York, NY, United States |
| Melissa Miller | Pfizer, New York, NY, United States |
| Xinli Hu | Pfizer, New York, NY, United States |
| Linda McCarthy | GlaxoSmithKline, Brentford, United Kingdom |

**Rheumatology Group**

| | |
|---|---|
| Kari Eklund | Hospital District of Helsinki and Uusimaa, Helsinki, Finland |
| Antti Palomäki | Hospital District of Southwest Finland, Turku, Finland |
| Pia Isomäki | Pirkanmaa Hospital District, Tampere, Finland |
| Laura Pirilä | Hospital District of Southwest Finland, Turku, Finland |
| Oili Kaipiainen-Seppänen | Northern Savo Hospital District, Kuopio, Finland |
| Johanna Huhtakangas | Northern Ostrobothnia Hospital District, Oulu, Finland |
| Bob Georgantas | Abbvie, Chicago, IL, United States |
| Jeffrey Waring | Abbvie, Chicago, IL, United States |
| Fedik Rahimov | Abbvie, Chicago, IL, United States |
| Apinya Lertratanakul | Abbvie, Chicago, IL, United States |
| Nizar Smaoui | Abbvie, Chicago, IL, United States |
| Anne Lehtonen | Abbvie, Chicago, IL, United States |
| David Close | Astra Zeneca, Cambridge, United Kingdom |
| Marla Hochfeld | Celgene, Summit, NJ, United States |
| Natalie Bowers | Genentech, San Francisco, CA, United States |
| John Michon | Genentech, San Francisco, CA, United States |
| Dorothee Diogo | Merck, Kenilworth, NJ, United States |
| Vinay Mehta | Merck, Kenilworth, NJ, United States |
| Kirsi Kalpala | Pfizer, New York, NY, United States |
| Nan Bing | Pfizer, New York, NY, United States |
| Xinli Hu | Pfizer, New York, NY, United States |
| Jorge Esparza Gordillo | GlaxoSmithKline, Brentford, United Kingdom |
| Nina Mars | Institute for Molecular Medicine Finland, HiLIFE, Helsinki, Finland |

**Pulmonology Group**

| | |
|---|---|
| Tarja Laitinen | Pirkanmaa Hospital District, Tampere, Finland |
| Margit Pelkonen | Northern Savo Hospital District, Kuopio, Finland |
| Paula Kauppi | Hospital District of Helsinki and Uusimaa, Helsinki, Finland |
| Hannu Kankaanranta | Pirkanmaa Hospital District, Tampere, Finland |
| Terttu Harju | Northern Ostrobothnia Hospital District, Oulu, Finland |
| Nizar Smaoui | Abbvie, Chicago, IL, United States |
| David Close | Astra Zeneca, Cambridge, United Kingdom |
| Susan Eaton | Biogen, Cambridge, MA, United States |
| Steven Greenberg | Celgene, Summit, NJ, United States |
| Hubert Chen | Genentech, San Francisco, CA, United States |
| Natalie Bowers | Genentech, San Francisco, CA, United States |
| John Michon | Genentech, San Francisco, CA, United States |
| Vinay Mehta | Merck, Kenilworth, NJ, United States |
| Jo Betts | GlaxoSmithKline, Brentford, United Kingdom |
| Soumitra Ghosh | GlaxoSmithKline, Brentford, United Kingdom |

**Cardiometabolic Diseases Group**

| | |
|---|---|
| Veikko Salomaa | The National Institute of Health and Welfare Helsinki, Finland |
| Teemu Niiranen | The National Institute of Health and Welfare Helsinki, Finland |
| Markus Juonala | Hospital District of Southwest Finland, Turku, Finland |
| Kaj Metsärinne | Hospital District of Southwest Finland, Turku, Finland |
| Mika Kähönen | Pirkanmaa Hospital District, Tampere, Finland |
| Juhani Junttila | Northern Ostrobothnia Hospital District, Oulu, Finland |
| Markku Laakso | Northern Savo Hospital District, Kuopio, Finland |
| Jussi Pihlajamäki | Northern Savo Hospital District, Kuopio, Finland |
| Juha Sinisalo | Hospital District of Helsinki and Uusimaa, Helsinki, Finland |
| Marja-Riitta Taskinen | Hospital District of Helsinki and Uusimaa, Helsinki, Finland |
| Tiinamaija Tuomi | Hospital District of Helsinki and Uusimaa, Helsinki, Finland |
| Jari Laukkanen | Central Finland Health Care District, Jyväskylä, Finland |

| | |
|---|---|
| Ben Challis | Astra Zeneca, Cambridge, United Kingdom |
| Andrew Peterson | Genentech, San Francisco, CA, United States |
| Julie Hunkapiller | Genentech, San Francisco, CA, United States |
| Natalie Bowers | Genentech, San Francisco, CA, United States |
| John Michon | Genentech, San Francisco, CA, United States |
| Dorothee Diogo | Merck, Kenilworth, NJ, United States |
| Audrey Chu | Merck, Kenilworth, NJ, United States |
| Vinay Mehta | Merck, Kenilworth, NJ, United States |
| Jaakko Parkkinen | Pfizer, New York, NY, United States |
| Melissa Miller | Pfizer, New York, NY, United States |
| Anthony Muslin | Sanofi, Paris, France |
| Dawn Waterworth | GlaxoSmithKline, Brentford, United Kingdom |

**Oncology Group**

| | |
|---|---|
| Heikki Joensuu | Hospital District of Helsinki and Uusimaa, Helsinki, Finland |
| Olli Carpén | Hospital District of Helsinki and Uusimaa, Helsinki, Finland |
| Tuomo Meretoja | Hospital District of Helsinki and Uusimaa, Helsinki, Finland |
| Lauri Aaltonen | Hospital District of Helsinki and Uusimaa, Helsinki, Finland |
| Johanna Mattson | Hospital District of Helsinki and Uusimaa, Helsinki, Finland |
| Johanna Schleutker | University of Turku, Turku, Finland |
| Annika Auranen | Pirkanmaa Hospital District , Tampere, Finland |
| Peeter Karihtala | Northern Ostrobothnia Hospital District, Oulu, Finland |
| Saila Kauppila | Northern Ostrobothnia Hospital District, Oulu, Finland |
| Päivi Auvinen | Northern Savo Hospital District, Kuopio, Finland |
| Klaus Elenius | Hospital District of Southwest Finland, Turku, Finland |
| Relja Popovic | Abbvie, Chicago, IL, United States |
| Jeffrey Waring | Abbvie, Chicago, IL, United States |
| Bridget Riley-Gillis | Abbvie, Chicago, IL, United States |
| Anne Lehtonen | Abbvie, Chicago, IL, United States |
| Athena Matakidou | Astra Zeneca, Cambridge, United Kingdom |
| Jennifer Schutzman | Genentech, San Francisco, CA, United States |
| Julie Hunkapiller | Genentech, San Francisco, CA, United States |
| Natalie Bowers | Genentech, San Francisco, CA, United States |
| John Michon | Genentech, San Francisco, CA, United States |
| Vinay Mehta | Merck, Kenilworth, NJ, United States |
| Andrey Loboda | Merck, Kenilworth, NJ, United States |
| Aparna Chhibber | Merck, Kenilworth, NJ, United States |
| Heli Lehtonen | Pfizer, New York, NY, United States |
| Stefan McDonough | Pfizer, New York, NY, United States |
| Marika Crohns | Sanofi, Paris, France |
| Diptee Kulkarni | GlaxoSmithKline, Brentford, United Kingdom¨ |

**Opthalmology Group**

| | |
|---|---|
| Kai Kaarniranta | Northern Savo Hospital District, Kuopio, Finland |
| Joni A Turunen | Hospital District of Helsinki and Uusimaa, Helsinki, Finland |
| Terhi Ollila | Hospital District of Helsinki and Uusimaa, Helsinki, Finland |
| Sanna Seitsonen | Hospital District of Helsinki and Uusimaa, Helsinki, Finland |
| Hannu Uusitalo | Pirkanmaa Hospital District, Tampere, Finland |
| Vesa Aaltonen | Hospital District of Southwest Finland, Turku, Finland |
| Hannele Uusitalo-Järvinen | Pirkanmaa Hospital District, Tampere, Finland |
| Marja Luodonpää | Northern Ostrobothnia Hospital District, Oulu, Finland |
| Nina Hautala | Northern Ostrobothnia Hospital District, Oulu, Finland |
| Heiko Runz | Biogen, Cambridge, MA, United States |
| Stephanie Loomis | Biogen, Cambridge, MA, United States |
| Erich Strauss | Genentech, San Francisco, CA, United States |
| Natalie Bowers | Genentech, San Francisco, CA, United States |

| | |
|---|---|
| Hao Chen | Genentech, San Francisco, CA, United States |
| John Michon | Genentech, San Francisco, CA, United States |
| Anna Podgornaia | Merck, Kenilworth, NJ, United States |
| Vinay Mehta | Merck, Kenilworth, NJ, United States |
| Dorothee Diogo | Merck, Kenilworth, NJ, United States |
| Joshua Hoffman | GlaxoSmithKline, Brentford, United Kingdom |

**Dermatology Group**

| | |
|---|---|
| Kaisa Tasanen | Northern Ostrobothnia Hospital District, Oulu, Finland |
| Laura Huilaja | Northern Ostrobothnia Hospital District, Oulu, Finland |
| Katariina Hannula-Jouppi | Hospital District of Helsinki and Uusimaa, Helsinki, Finland |
| Teea Salmi | Pirkanmaa Hospital District, Tampere, Finland |
| Sirkku Peltonen | Hospital District of Southwest Finland, Turku, Finland |
| Leena Koulu | Hospital District of Southwest Finland, Turku, Finland |
| Ilkka Harvima | Northern Savo Hospital District, Kuopio, Finland |
| Kirsi Kalpala | Pfizer, New York, NY, United States |
| Ying Wu | Pfizer, New York, NY, United States |
| David Choy | Genentech, San Francisco, CA, United States |
| John Michon | Genentech, San Francisco, CA, United States |
| Nizar Smaoui | Abbvie, Chicago, IL, United States |
| Fedik Rahimov | Abbvie, Chicago, IL, United States |
| Anne Lehtonen | Abbvie, Chicago, IL, United States |
| Dawn Waterworth | GlaxoSmithKline, Brentford, United Kingdom |

**Odontology Group**

| | |
|---|---|
| Pirkko Pussinen | Hospital District of Helsinki and Uusimaa, Helsinki, Finland |
| Aino Salminen | Hospital District of Helsinki and Uusimaa, Helsinki, Finland |
| Tuula Salo | Hospital District of Helsinki and Uusimaa, Helsinki, Finland |
| David Rice | Hospital District of Helsinki and Uusimaa, Helsinki, Finland |
| Pekka Nieminen | Hospital District of Helsinki and Uusimaa, Helsinki, Finland |
| Ulla Palotie | Hospital District of Helsinki and Uusimaa, Helsinki, Finland |
| Maria Siponen | Northern Savo Hospital District, Kuopio, Finland |
| Liisa Suominen | Northern Savo Hospital District, Kuopio, Finland |
| Päivi Mäntylä | Northern Savo Hospital District, Kuopio, Finland |
| Ulvi Gursoy | Hospital District of Southwest Finland, Turku, Finland |
| Vuokko Anttonen | Northern Ostrobothnia Hospital District, Oulu, Finland |
| Kirsi Sipilä | Northern Ostrobothnia Hospital District, Oulu, Finland |

#### FinnGen Analysis working group

| | |
|---|---|
| Justin Wade Davis | Abbvie, Chicago, IL, United States |
| Bridget Riley-Gillis | Abbvie, Chicago, IL, United States |
| Danjuma Quarless | Abbvie, Chicago, IL, United States |
| Fedik Rahimov | Abbvie, Chicago, IL, United States |
| Sahar Esmaeeli | Abbvie, Chicago, IL, United States |
| Slavé Petrovski | Astra Zeneca, Cambridge, United Kingdom |
| Eleonor Wigmore | Astra Zeneca, Cambridge, United Kingdom |
| Jimmy Liu | Biogen, Cambridge, MA, United States |
| Chia-Yen Chen | Biogen, Cambridge, MA, United States |
| Paola Bronson | Biogen, Cambridge, MA, United States |
| Ellen Tsai | Biogen, Cambridge, MA, United States |
| Stephanie Loomis | Biogen, Cambridge, MA, United States |
| Yunfeng Huang | Biogen, Cambridge, MA, United States |
| Joseph Maranville | Celgene, Summit, NJ, United States |
| Shameek Biswas | Celgene, Summit, NJ, United States |
| Elmutaz SE Mohammed | Celgene, Summit, NJ, United States |
| Samir Wadhawan | Bristol-Meyers-Squibb |

| | |
|---|---|
| Erika Kvikstad | Bristol-Meyers-Squibb |
| Minal Caliskan | Bristol-Meyers-Squibb |
| Diana Chang | Genentech, San Francisco, CA, United States |
| Julie Hunkapiller | Genentech, San Francisco, CA, United States |
| Tushar Bhangale | Genentech, San Francisco, CA, United States |
| Natalie Bowers | Genentech, San Francisco, CA, United States |
| Sarah Pendergrass | Genentech, San Francisco, CA, United States |
| Dorothee Diogo | Merck, Kenilworth, NJ, United States |
| Emily Holzinger | Merck, Kenilworth, NJ, United States |
| Padhraig Gormley | Merck, Kenilworth, NJ, United States |
| Xing Chen | Pfizer, New York, NY, United States |
| Åsa Hedman | Pfizer, New York, NY, United States |
| Karen S King | GlaxoSmithKline, Brentford, United Kingdom |
| Clarence Wang | Sanofi, Paris, France |
| Ethan Xu | Sanofi, Paris, France |
| Franck Auge | Sanofi, Paris, France |
| Clement Chatelain | Sanofi, Paris, France |
| Deepak Rajpal | Sanofi, Paris, France |
| Dongyu Liu | Sanofi, Paris, France |
| Katherine Call | Sanofi, Paris, France |
| Tai-he Xia | Sanofi, Paris, France |
| Beryl Cummings | Maze Therapeutics, San Francisco, CA, United States |
| Matt Brauer | Maze Therapeutics, San Francisco, CA, United States |
| Mitja Kurki | Institute for Molecular Medicine Finland, HiLIFE, University of Helsinki, Finland / Broad Institute, Cambridge, MA, United States |
| Samuli Ripatti | Institute for Molecular Medicine Finland, HiLIFE, University of Helsinki, Finland |
| Mark Daly | Institute for Molecular Medicine Finland, HiLIFE, University of Helsinki, Finland |
| Juha Karjalainen | Institute for Molecular Medicine Finland, HiLIFE, University of Helsinki, Finland / Broad Institute, Cambridge, MA, United States |
| Aki Havulinna | Institute for Molecular Medicine Finland, HiLIFE, University of Helsinki, Finland |
| Anu Jalanko | Institute for Molecular Medicine Finland, HiLIFE, University of Helsinki, Finland |
| Priit Palta | Institute for Molecular Medicine Finland, HiLIFE, University of Helsinki, Finland |
| Pietro della B Parolo | Institute for Molecular Medicine Finland, HiLIFE, University of Helsinki, Finland |
| Wei Zhou | Broad Institute, Cambridge, MA, United States |
| Susanna Lemmelä | Institute for Molecular Medicine Finland, HiLIFE, University of Helsinki, Finland |
| Manuel Rivas | University of Stanford, Stanford, CA, United States |
| Jarmo Harju | Institute for Molecular Medicine Finland, HiLIFE, University of Helsinki, Finland |
| Aarno Palotie | Institute for Molecular Medicine Finland, HiLIFE, University of Helsinki, Finland |
| Arto Lehisto | Institute for Molecular Medicine Finland, HiLIFE, University of Helsinki, Finland |
| Andrea Ganna | Institute for Molecular Medicine Finland, HiLIFE, University of Helsinki, Finland |
| Vincent Llorens | Institute for Molecular Medicine Finland, HiLIFE, University of Helsinki, Finland |
| Hannele Laivuori | Institute for Molecular Medicine Finland, HiLIFE, University of Helsinki, Finland |
| Sina Rüeger | Institute for Molecular Medicine Finland, HiLIFE, University of Helsinki, Finland |
| Mari E Niemi | Institute for Molecular Medicine Finland, HiLIFE, University of Helsinki, Finland |
| Taru Tukiainen | Institute for Molecular Medicine Finland, HiLIFE, University of Helsinki, Finland |
| Mary Pat Reeve | Institute for Molecular Medicine Finland, HiLIFE, University of Helsinki, Finland |
| Henrike Heyne | Institute for Molecular Medicine Finland, HiLIFE, University of Helsinki, Finland |
| Nina Mars | Institute for Molecular Medicine Finland, HiLIFE, University of Helsinki, Finland |
| Kimmo Palin | University of Helsinki, Helsinki, Finland |
| Javier Garcia-Tabuenca | University of Tampere, Tampere, Finland |
| Harri Siirtola | University of Tampere, Tampere, Finland |
| Tuomo Kiiskinen | Institute for Molecular Medicine Finland, HiLIFE, University of Helsinki, Finland |
| Jiwoo Lee | Institute for Molecular Medicine Finland, HiLIFE, University of Helsinki, Finland / Broad Institute, Cambridge, MA, United States |
| Kristin Tsuo | Institute for Molecular Medicine Finland, HiLIFE, University of Helsinki, Finland / Broad Institute, Cambridge, MA, United States |

Amanda Elliott Institute for Molecular Medicine Finland, HiLIFE, University of Helsinki, Finland / Broad Institute, Cambridge, MA, United States
Kati Kristiansson THL Biobank / The National Institute of Health and Welfare Helsinki, Finland
Mikko Arvas Finnish Red Cross Blood Service, Helsinki, Finland
Kati Hyvärinen Finnish Red Cross Blood Service, Helsinki, Finland
Jarmo Ritari Finnish Red Cross Blood Service, Helsinki, Finland
Miika Koskinen Helsinki Biobank / Helsinki University and Hospital District of Helsinki and Uusimaa, Helsinki
Olli Carpén Helsinki Biobank / Helsinki University and Hospital District of Helsinki and Uusimaa, Helsinki
Johannes Kettunen Northern Finland Biobank Borealis / University of Oulu / Northern Ostrobothnia Hospital District, Oulu, Finland
Katri Pylkäs Northern Finland Biobank Borealis / University of Oulu / Northern Ostrobothnia Hospital District, Oulu, Finland
Marita Kalaoja Northern Finland Biobank Borealis / University of Oulu / Northern Ostrobothnia Hospital District, Oulu, Finland
Minna Karjalainen Northern Finland Biobank Borealis / University of Oulu / Northern Ostrobothnia Hospital District, Oulu, Finland
Tuomo Mantere Northern Finland Biobank Borealis / University of Oulu / Northern Ostrobothnia Hospital District, Oulu, Finland
Eeva Kangasniemi Finnish Clinical Biobank Tampere / University of Tampere / Pirkanmaa Hospital District, Tampere, Finland
Sami Heikkinen Biobank of Eastern Finland / University of Eastern Finland / Northern Savo Hospital District, Kuopio, Finland
Arto Mannermaa Biobank of Eastern Finland / University of Eastern Finland / Northern Savo Hospital District, Kuopio, Finland
Eija Laakkonen Central Finland Biobank / University of Jyväskylä / Central Finland Health Care District, Jyväskylä, Finland
Csilla Sipeky University of Turku, Turku, Finland
Samuel Heron University of Turku, Turku, Finland
Antti Karlsson Auria Biobank / University of Turku / Hospital District of Southwest Finland, Turku, Finland
Dhanaprakash Jambulingam University of Turku, Turku, Finland
Venkat Subramaniam Rathinakannan University of Turku, Turku, Finland

#### Biobank directors

Lila Kallio Auria Biobank / University of Turku / Hospital District of Southwest Finland, Turku, Finland
Sirpa Soini THL Biobank / The National Institute of Health and Welfare Helsinki, Finland
Jukka Partanen Finnish Red Cross Blood Service / Finnish Hematology Registry and Clinical Biobank, Helsinki, Finland
Eero Punkka Helsinki Biobank / Helsinki University and Hospital District of Helsinki and Uusimaa, Helsinki
Raisa Serpi Northern Finland Biobank Borealis / University of Oulu / Northern Ostrobothnia Hospital District, Oulu, Finland
Johanna Mäkelä Finnish Clinical Biobank Tampere / University of Tampere / Pirkanmaa Hospital District, Tampere, Finland
Veli-Matti Kosma Biobank of Eastern Finland / University of Eastern Finland / Northern Savo Hospital District, Kuopio, Finland
Teijo Kuopio Central Finland Biobank / University of Jyväskylä / Central Finland Health Care District, Jyväskylä, Finland

#### FinnGen Teams

**Administration**

Anu Jalanko Institute for Molecular Medicine Finland, HiLIFE, University of Helsinki, Finland
Risto Kajanne Institute for Molecular Medicine Finland, HiLIFE, University of Helsinki, Finland

Mervi Aavikko Institute for Molecular Medicine Finland, HiLIFE, University of Helsinki, Finland
Manuel González Jiménez Institute for Molecular Medicine Finland, HiLIFE, University of Helsinki, Finland
**Analysis**
Mitja Kurki Institute for Molecular Medicine Finland, HiLIFE, University of Helsinki, Finland / Broad Institute, Cambridge, MA, United States
Juha Karjalainen Institute for Molecular Medicine Finland, HiLIFE, University of Helsinki, Finland / Broad Institute, Cambridge, MA, United States
Pietro della B Parola Institute for Molecular Medicine Finland, HiLIFE, University of Helsinki, Finland
Sina Rüeger Institute for Molecular Medicine Finland, HiLIFE, University of Helsinki, Finland
Arto Lehistö Institute for Molecular Medicine Finland, HiLIFE, University of Helsinki, Finland
Wei Zhou Broad Institute, Cambridge, MA, United States
Masahiro Kanai Broad Institute, Cambridge, MA, United States
**Clinical Endpoint Development**
Hannele Laivuori Institute for Molecular Medicine Finland, HiLIFE, University of Helsinki, Finland
Aki Havulinna Institute for Molecular Medicine Finland, HiLIFE, University of Helsinki, Finland
Susanna Lemmelä Institute for Molecular Medicine Finland, HiLIFE, University of Helsinki, Finland
Tuomo Kiiskinen Institute for Molecular Medicine Finland, HiLIFE, University of Helsinki, Finland
**Communication**
Mari Kaunisto Institute for Molecular Medicine Finland, HiLIFE, University of Helsinki, Finland
**Data Management and IT Infrastructure**
Jarmo Harju Institute for Molecular Medicine Finland, HiLIFE, University of Helsinki, Finland
Elina Kilpeläinen Institute for Molecular Medicine Finland, HiLIFE, University of Helsinki, Finland
Timo P. Sipilä Institute for Molecular Medicine Finland, HiLIFE, University of Helsinki, Finland
Georg Brein Institute for Molecular Medicine Finland, HiLIFE, University of Helsinki, Finland
Oluwaseun A. Dada Institute for Molecular Medicine Finland, HiLIFE, University of Helsinki, Finland
Ghazal Awaisa Institute for Molecular Medicine Finland, HiLIFE, University of Helsinki, Finland
Anastasia Shcherban Institute for Molecular Medicine Finland, HiLIFE, University of Helsinki, Finland
**Genotyping**
Kati Donner Institute for Molecular Medicine Finland, HiLIFE, University of Helsinki, Finland
Timo P. Sipilä Institute for Molecular Medicine Finland, HiLIFE, University of Helsinki, Finland
**Sample Collection Coordination**
Anu Loukola Helsinki Biobank / Helsinki University and Hospital District of Helsinki and Uusimaa, Helsinki
**Sample Logistics**
Päivi Laiho THL Biobank / The National Institute of Health and Welfare Helsinki, Finland
Tuuli Sistonen THL Biobank / The National Institute of Health and Welfare Helsinki, Finland
Essi Kaiharju THL Biobank / The National Institute of Health and Welfare Helsinki, Finland
Markku Laukkanen THL Biobank / The National Institute of Health and Welfare Helsinki, Finland
Elina Järvensivu THL Biobank / The National Institute of Health and Welfare Helsinki, Finland
Sini Lähteenmäki THL Biobank / The National Institute of Health and Welfare Helsinki, Finland
Lotta Männikkö THL Biobank / The National Institute of Health and Welfare Helsinki, Finland
Regis Wong THL Biobank / The National Institute of Health and Welfare Helsinki, Finland
**Registry Data Operations**
Hannele Mattsson THL Biobank / The National Institute of Health and Welfare Helsinki, Finland
Kati Kristiansson THL Biobank / The National Institute of Health and Welfare Helsinki, Finland
Susanna Lemmelä Institute for Molecular Medicine Finland, HiLIFE, University of Helsinki, Finland
Tero Hiekkalinna THL Biobank / The National Institute of Health and Welfare Helsinki, Finland
Teemu Paajanen THL Biobank / The National Institute of Health and Welfare Helsinki, Finland
**Sequencing Informatics**
Priit Palta Institute for Molecular Medicine Finland, HiLIFE, University of Helsinki, Finland
Kalle Pärn Institute for Molecular Medicine Finland, HiLIFE, University of Helsinki, Finland
**Trajectory Team**
Tarja Laitinen Pirkanmaa Hospital District, Tampere, Finland
Harri Siirtola University of Tampere, Tampere, Finland
Javier Gracia-Tabuenca University of Tampere, Tampere, Finland
